# The MAP She1 coordinates directional spindle positioning by spatially restricting dynein activity

**DOI:** 10.1101/2021.02.02.429374

**Authors:** Kari H. Ecklund, Megan E. Bailey, Carsten K. Dietvorst, Charles L. Asbury, Steven M. Markus

## Abstract

Dynein motors move the mitotic spindle to the cell division plane in many cell types, including in budding yeast, in which dynein is assisted by numerous factors including the microtubule-associated protein (MAP) She1. Evidence suggests that She1 plays a role in polarizing dynein-mediated spindle movements toward the daughter cell; however, how She1 performs this function is unknown. We find that She1 assists dynein in maintaining the spindle close to the bud neck, such that at anaphase onset the chromosomes are segregated to mother and daughter cells. She1 does so by attenuating the initiation of dynein-mediated spindle movements specifically within the mother cell, ensuring such movements are polarized toward the daughter cell. Our data indicate that this activity relies on She1 binding to the microtubule-bound conformation of the dynein microtubule-binding domain, and to astral microtubules within mother cells. Our findings reveal how an asymmetrically localized MAP directionally tunes dynein activity by attenuating motor activity in a spatially confined manner.

## INTRODUCTION

By transporting various cargoes along microtubules, the dynein and kinesin families of molecular motors play important roles in many cellular processes, including coordinating the spatially and temporally appropriate positions of membrane-bound vesicles, organelles, and the mitotic spindle. In addition to being affected by various motor-specific accessories and regulators, microtubule motors are regulated by a family of <u>M</u>icrotubule-<u>A</u>ssociated <u>P</u>roteins (MAPs). For instance, the neuronal MAP tau, well known for its implication in Alzheimer’s disease^1^, inhibits kinesin-1-based transport of vesicles and organelles in neurons *in vivo*, and induces detachment and pausing behavior of kinesin-1 and dynein *in vitro*^2–5^. In addition to tau, several other MAPs, including MAP4 and MAP9 affect dynein transport functions *in vivo* and *in vitro*^6–8^. Although the mechanisms by which MAPs affect *in vitro* motor motility are beginning to be understood^5,7,9,10^, how their activities result in the appropriate positioning of the various dynein and kinesin cargoes in cells is unclear.

She1 (<u>S</u>ensitive to <u>h</u>igh expression-<u>1</u>) is a yeast-specific MAP that has been shown to potently affect dynein motility *in vitro*, and dynein-mediated spindle positioning *in vivo*^11–13^. Although the precise mechanism by which She1 promotes appropriate *in vivo* dynein activity is unclear, our recent *in vitro* data determined that She1 affects dynein motility through simultaneous interactions with the dynein microtubule binding domain (MTBD) and the microtubule^10^. In addition to reducing dynein velocity, She1 reduces the dynein-microtubule dissociation rate^10^, suggesting that it may promote dynein-microtubule interaction in cells, and potentially affect dynein force production. Given that dynein functions in concert with the cortical receptor Num1 (potentially orthologous to NuMA in humans)^14^ and the dynactin complex (<u>dyne</u>in <u>act</u>ivator)^15–18^ in cells, it is unclear whether these prior findings with dynein alone apply to active Num1-dynein-dynactin complexes in cells. Thus, a clear picture of how this MAP affects dynein-mediated spindle positioning is lacking.

She1 is one of several known effectors of budding yeast dynein activity, which also include Ndl1 (Nde1 in humans)^19^, Bik1 (CLIP170 in humans)^20^, and Pac1 (LIS1 in humans)^16^. In contrast to higher eukaryotes, the only known function for dynein in this organism is to position the nucleus with the enclosed mitotic spindle at the future site of cytokinesis^21,22^: the narrow neck between the mother and daughter cells. Dynein performs this activity from cortical Num1 receptor sites, to which it is delivered by a multi-step ‘offloading’ mechanism^16,20,23,24^. Specifically: (1) dynein indirectly associates with the plus ends of dynamic microtubules in a Pac1 and Bik1-dependent manner^16,20,25,26^; (2) the dynactin complex is recruited to plus end-bound dynein^17^; (3) upon encountering cortical Num1 receptors, dynein and dynactin are offloaded and activated to translocate the nucleus and spindle^24,25^. Although the offloading process appears to be biased towards the daughter cell^24^, a greater number of dynein foci are apparent in mother cells, likely due to their age-dependent accumulation. In spite of this imbalance, dynein-mediated spindle movements toward the daughter cell prevail. How dynein activity is appropriately biased to ensure proper spindle position is unclear.

Our recent studies discovered that She1 is a key effector that polarizes dynein-mediated spindle positioning. Specifically, we found that deletion of She1 compromises dynein-mediated spindle translocation events that lead to the spindle crossing the bud neck, and results in a higher prevalence of anaphase onset within the mother cells, quite distal from the bud neck^13^. How She1 performs this function, however, is unknown. In this study, we employ a combination of *in vitro* and *in vivo* methods to determine the precise basis by which She1 affects dynein activity. In particular, we find that our previous *in vitro* data indicating a role for She1 in reducing dynein velocity and promoting dynein-microtubule binding do not reflect the likely basis for in-cell activity. Rather, we find that She1 supports dynein-mediated spindle positioning by attenuating the initiation of cortical dynein-dynactin-mediated spindle movements predominantly in the mother cell. We find that this activity requires She1 interactions with the microtubule-bound conformation of the dynein microtubule-binding domain as well as microtubules, partially reconciling our *in vivo* and *in vitro* data. Finally, we find that the likely basis for mother cell-specific inhibition of dynein activity is the asymmetric binding of She1 to astral microtubules within this compartment. In summary, our findings describe how an asymmetrically localized MAP can spatially tune the activity of a molecular motor to coordinate appropriate cargo transport.

## RESULTS

### She1 is required for normal spindle positioning and mitotic timing

We previously found that She1 is important for polarizing dynein-mediated spindle movements towards the daughter cell^13^. Consistent with its importance in spindle positioning, single time-point images acquired of cells grown at low temperatures (16°C; to exacerbate dynamic microtubule-mediated processes) revealed a mild spindle positioning defect^10^. To gain additional insight into the role of She1 in dynein-mediated spindle positioning and cell cycle progression, we performed time lapse imaging of cells over the course of several cell cycles using a microfluidics-based platform (CellAsic ONIX, Millipore Sigma), in which cells are imaged at 30°C in the presence of constantly replenished nutrients. We imaged wild-type (*SHE1*) and *she1*Δ cells expressing Spc110-Venus, NLS-3mCherry, and mTurquoise2-Tub1, to image spindle pole bodies (SPBs), the nucleus, and microtubules, respectively. Consistent with the low temperature single time-point assay^10^, 17.7% of *she1*Δ cells (compared to only 2.5% of *SHE1* cells) exhibited mispositioned spindles at anaphase onset, confirming the importance of She1 in normal spindle positioning (Fig. 1A and B).

**Figure 1.**
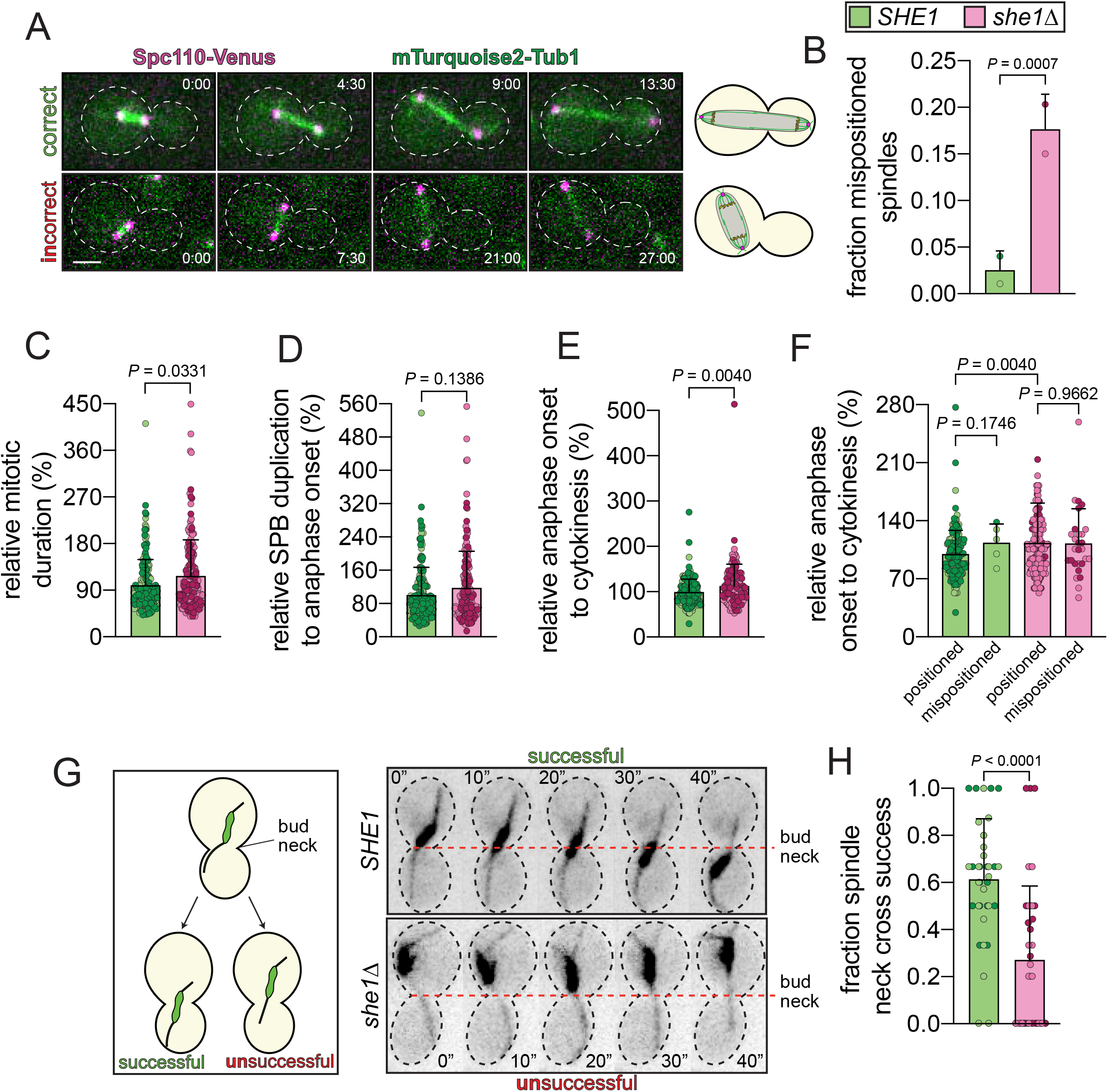
She1 is required for normal spindle position and cell cycle progression. (A) Representative time-lapse images of cells expressing NLS-3mCherry (not shown here; see Fig. S1), Spc110-Venus, and mTurquoise2-Tub1 depicting spindle position (correct and incorrect) at the moments preceding and during onset of anaphase (cartoon on the right depicts the last frame of each respective time series; minutes and seconds are indicated). To control for imaging conditions (room temperature in particular), wild-type and *she1*Δ cells were imaged together (for each independent replicate; the motorized stage on the microscope was used to switch between chambers) by introducing respective yeast strains into two different chambers of the CellAsic ONIX system. (B) Plot depicting the mean fraction of mispositioned spindles in *SHE1* (wild-type) and *she1*Δ strains. Each diamond represents the mean fraction of mispositioned spindles from each independent replicate. Error bars represent standard deviation. P value was calculated from Z-score, as previously described^76^. (C - F) Plots depicting relative time intervals between various temporal landmarks (see Fig. S1). Data were normalized [*i* = (*x*-minimum value from control)/(maximum value from control-minimum value from control), where *i* is the normalized value, and *x* is the nonnormalized value] to account for differences between independent replicates due to differences in imaging conditions (likely due to fluctuations in room temperature). See Figure S1 for raw data (for panels B – F, n = 191 cells for wild-type, and n = 163 cells for *she1*Δ cells, from 2 independent replicates each). P values were calculated using Mann-Whitney tests. (G) Representative inverse fluorescence time-lapse images from hydroxyurea-arrested *SHE1 kar9*Δ (top) and *she1Δ kar9*Δ (bottom) cells expressing GFP-Tub1 (along with cartoon schematic) depicting a successful (top) or unsuccessful (bottom) attempt of the spindle to cross the bud neck subsequent to the initiation of dynein-mediated spindle movement toward the neck (neck indicated by dashed red line). Dynein-mediated spindle movements were identified as such by the apparent directed migration of the spindle following and coincident with an astral microtubule associating with the cell cortex. (H) Plot depicting the fraction of bud neck-directed dynein-mediated spindle movements that result in the successful migration of the spindle midpoint across the bud neck. Circles represent the fraction of successful spindle neck crosses per cell (n = 40 cells from 2 independent replicates for each, with n = 194 and 146 observed spindle neck cross attempts from *SHE1* and *she1*Δ cells, respectively). The P value was calculated using an unpaired two-tailed Welch’s t-test.

Given that She1 is important for proper spindle positioning, we wondered whether *she1*Δ cells exhibit any cell cycle delays. Cells with mispositioned spindles trigger a spindle positioning checkpoint (SPC), which delays cytokinesis and mitotic exit by preventing activation of the mitotic exit network (MEN)^27–31^. Of note, in addition to localizing to microtubules, She1 is also found at the bud neck, a site where various SPC effectors localize (*e.g*., Elm1, and Kin4^32–36^). We assessed several aspects of cell cycle progression by measuring the time intervals between various temporal landmarks (Fig. S1). This revealed an 11.0% increase in mitotic duration in *she1*Δ cells (Fig. 1C; defined as the duration between SPB duplication and spindle disassembly; timing was normalized to account for variability between independent replicates that were each internally controlled; see Fig. S1 and Methods) that was largely a consequence of a delay between anaphase onset and cytokinesis (Fig. 1E; 13.5%; *P* = 0.0040). This may be due to the delay previously noted in the onset of spindle disassembly in *she1*Δ cells^37^. We observed no significant delay between SPB duplication (a marker of initiation of spindle assembly) and anaphase onset due to She1 deletion (Fig. 1D), suggesting that the timing of spindle assembly is unperturbed upon She1 deletion. Separate analysis of cells with mispositioned spindles and those with properly positioned spindles revealed a 13.6% delay between anaphase onset and cytokinesis in *SHE1* cells (*P* = 0.1746; note the very small dataset due to the low prevalence of mispositioned spindles in wild-type cells; n = 5 cells, from 2 independent replicates), as would be expected given the presence of an intact SPC (Fig. 1F; as has been noted in numerous other studies^28–31^). However, *she1*Δ cells exhibited no such delay, suggesting that She1 may play a role in SPC function.

### She1 induces a persistent force generating dynein state in vitro but likely not in vivo

We sought to clarify the role of She1 in dynein-mediated spindle positioning. Our prior work found that She1 enhances dynein-microtubule binding affinity through simultaneous interactions with the dynein microtubule-binding domain (MTBD) and the microtubule^10^. We therefore hypothesized that She1 promotes daughter cell-directed dynein activity by enhancing its force generation capacity (by reducing dynein-microtubule dissociation). In this model, She1 assists dynein in generating sufficient force to pull the nucleus (with enclosed spindle) through the narrow mother-bud neck (which is approximately half the width of the nucleus; see Fig. 3A and B). If true, we predicted that loss of She1 would lead to scenarios where dynein-mediated spindle movements into the bud neck would be more prone to failure. To determine whether this is the case, we imaged dynein-mediated spindle movements in *SHE1* and *she1*Δ cells arrested in a metaphase-like state by treatment with the DNA-synthesis inhibitor hydroxyurea (HU). This arrest permits numerous observations of dynein-mediated spindle movements over a fairly short imaging period, with a large fraction of them involving translocation of the spindle across the bud neck. To ensure that all spindle movements are dynein dependent, we performed these and all similar subsequent experiments in cells deleted for *KAR9*, a key member of an actomyosin-mediated spindle orientation pathway^38^. Dynein-mediated spindle movements that were directed toward the bud neck were manually scored based on whether they resulted in a successful or unsuccessful crossing of the bud neck (Fig. 1G). Whereas 61.3% of such spindle movements resulted in a neck cross in wild-type cells, this value was reduced to 27.1% in *she1*Δ cells (Fig. 1H), lending support to an assisted force model for She1 function.

To determine if She1 affects dynein force generation, we employed optical trapping with a minimally processive, artificially dimerized dynein fragment that is affected by She1 in single molecule and ensemble motility assays^10,13^ (6His-GST-dynein_331_; adsorbed to polystyrene beads with an anti-6His antibody; Fig. 2A). Single dynein motor-coated beads (as determined by a bead-microtubule binding fraction < 0.3) captured in a fixed-position laser-based optical trap were lowered on to glass-immobilized microtubules, and their motility was measured with a high degree of spatial and temporal precision in the absence or presence of 2 or 5 nM recombinant She1-HaloTag^TMR^ (hereafter referred to as She1-TMR), concentrations previously determined to approximate cellular levels^13^. Consistent with our previous single molecule data, pre-stall bead velocity decreased significantly in the presence of increasing concentrations of She1-TMR (Fig. 2B and C). Although there was no appreciable change in dynein stall force in the presence of She1, we observed a 4.6-fold increase in stall time (from 20.2 to 92.7 seconds; Fig. 2B, D and E), indicating that She1 indeed induces a persistent force generating state for dynein *in vitro*.

**Figure 2.**
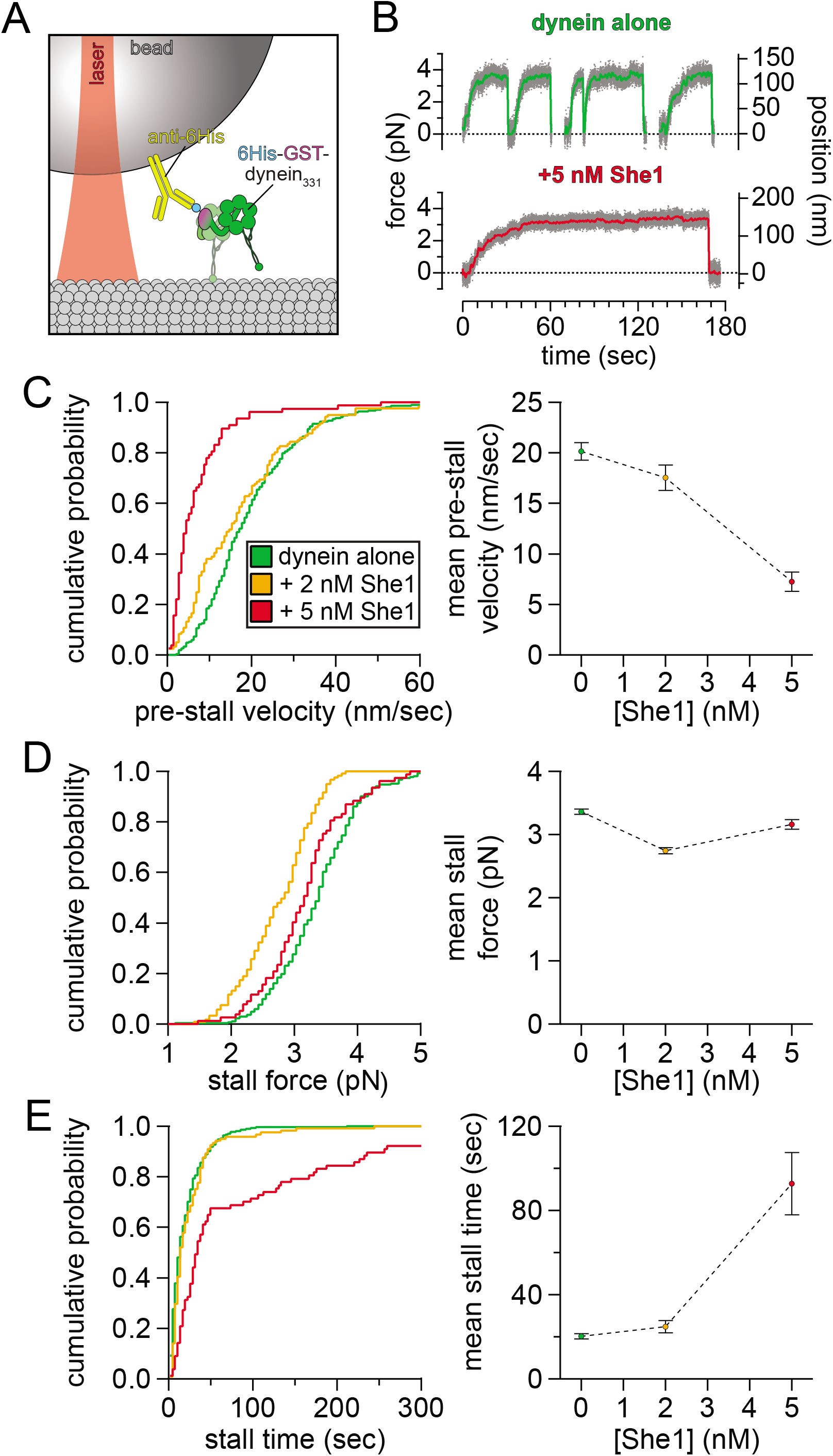
She1 increases dynein stall time but not stall force. (A and B) Schematic of optical trapping experimental setup (A) and representative traces (B). 6His-GST-dynein_331_-decorated polystyrene beads (attached using anti-6His antibody) lowered onto a coverslip-immobilized microtubule are transported away from the trap center until the motor can no longer overcome the opposing force of the trap (at which point it stalls and ultimately detaches). (C - E) Plots depicting cumulative probability frequencies (left) and mean values (right; ± standard error of the mean) of pre-stall velocity (C), stall force (D) and stall time (E) for polystyrene beads decorated with single molecules of 6His-GST-dynein_331_ (as determined from a microtubule-binding fraction of < 0.3; n = 272, 121, and 77 motility events recorded in the presence of 0, 2, and 5 nM She1-TMR, respectively).

Our results indicate that although She1 does not increase dynein force generation, it does improve the ability of dynein to remain bound to microtubules in the presence of an opposing force. In cells, this could potentially be provided by the large nucleus being squeezed through the narrow bud neck during dynein-mediated nuclear migration. To test this hypothesis *in vivo*, we again turned to live cell imaging, but this time with cells expressing Nup133-3mCherry (to visualize the nucleus) in addition to GFP-Tub1. HU-arrested *SHE1* and *she1*Δ cells (all of which were deleted for *KAR9*, as described above) were imaged, and the frequencies with which nuclei successfully crossed the neck were scored. In particular, those events that were directed toward the bud neck were manually scored based on whether they resulted in a successful or unsuccessful insertion of the nucleus into the narrow bud neck (see Fig. 3C). In addition to performing this assessment in otherwise wild-type cells, we also employed cells deleted for *BNI1*, a formin that when deleted results in a widening of the bud neck^39,40^ (from 1.1 μm to 1.5 μm) but not the nucleus (2.3 μm in both *BNI1* and *bni1*Δ Fig. 3A). We reasoned that if the bud neck provides a barrier over which dynein-mediated nuclear migration must overcome, and that She1 assists dynein in performing this function, then widening the bud neck would potentially rescue *she1*Δ phenotypes. Although we noted a small, but statistically insignificant increase in the success rate for dynein-mediated nuclear/neck translocation as a consequence of *BNI1* deletion, the presence or absence of She1 had no apparent effect on this phenomenon (*P* ≥ 0.8823; Fig. 3D). These data, which challenge our assisted force model, suggest that although the bud neck acts as a physical barrier to nuclear migration, She1 does not assist dynein in overcoming it. Thus, a model that reconciles our *in vitro* and *in vivo* data and accounts for She1 polarizing dynein mediated spindle movements toward the daughter cell remains lacking.

**Figure 3.**
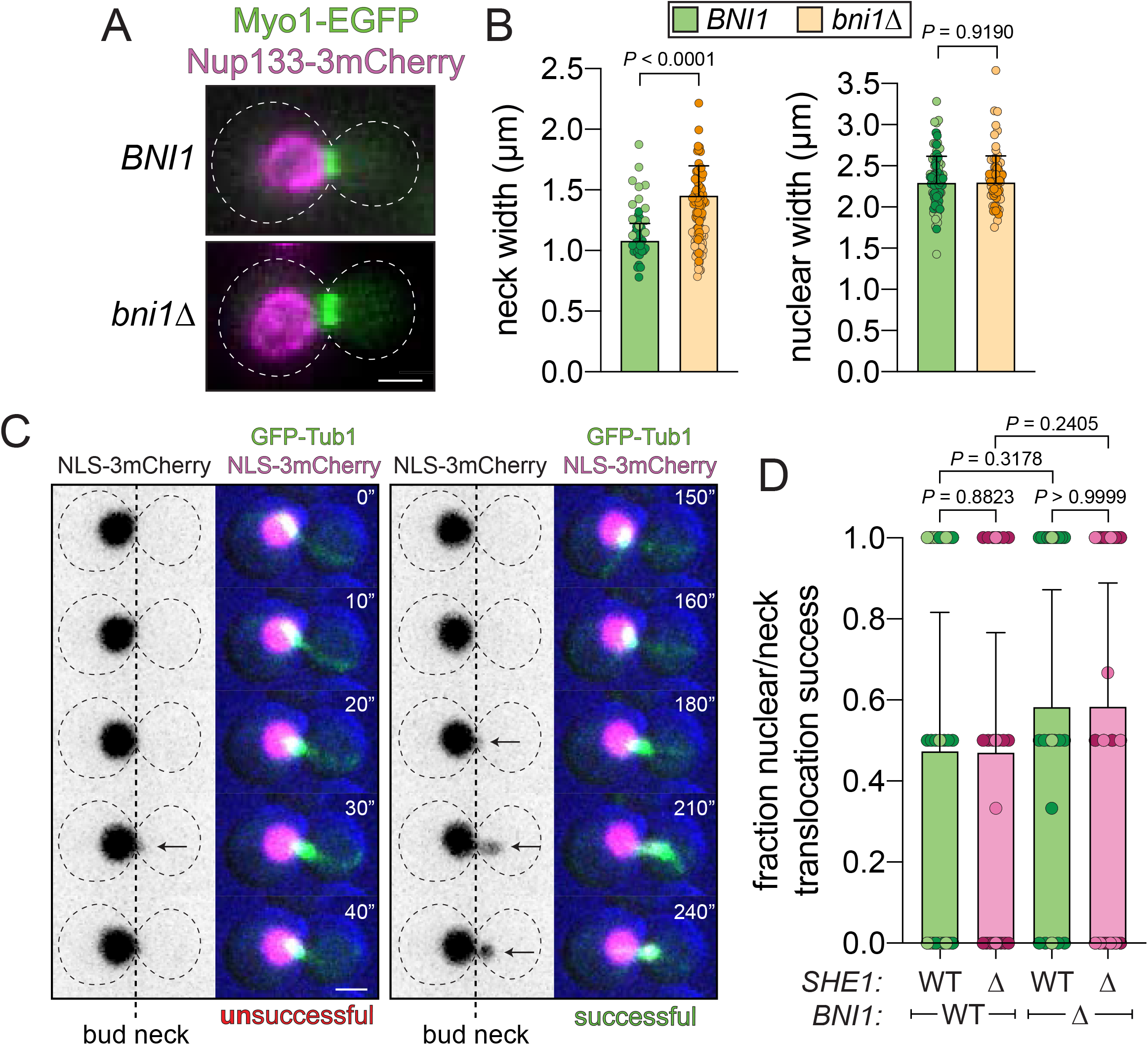
She1 has no impact on the ability of dynein to translocate the nucleus through the narrow bud neck. (A and B) Representative fluorescence images of hydroxyurea-arrested wild-type (*BNI1*) and *bni1*Δ cells expressing Nup133-3mCherry and Myo1-EFGP, and plots depicting measured width values of the bud neck (measured using Myo1-EGFP signal; n = 87 and 66 cells, from 2 independent replicates) and nucleus (measured at the widest point parallel to the axis of the bud neck; n = 88 and 67 cells, from 2 independent replicates). P values were calculated using an unpaired two-tailed Welch’s t-test (left) or a Mann-Whitney test (right). (C) Representative fluorescence time-lapse images of a hydroxyurea-arrested cell expressing NLS-3mCherry and GFP-Tub1. An example of an unsuccessful attempt to translocate the nucleus through the neck is shown on the left, while a successful attempt in the same cell is shown on the right. Arrows indicate the nucleus crossing the bud neck. Note that in the unsuccessful attempt (left), a dynein-mediated spindle migration event (apparent from microtubule-cortex association coupled with directed motion) leads to the nucleus crossing the neck for a short period before retracting into the mother cell (situated to the left of the daughter cell), whereas the nucleus remains within the neck following the subsequent successful attempt (right). (D) Plot depicting the fraction of events in which the nucleus successfully translocated the neck. We define a successful nuclear/neck translocation as an event in which the midpoint of the nucleus moves across the bud neck coincident with a dynein-mediated spindle movement. Circles represent the nuclear/neck translocation success rate per cell (from left to right, n = 33, 40, 43, and 42 cells, from 2 independent replicates; n = 65, 74, 96, and 88 nuclear translocation attempts were assessed from each). Error bars indicate standard deviation. P values were calculated using a Mann-Whitney test.

### She1 attenuates dynein activity in vivo and promotes bud neck proximal spindle position

Previous studies suggested that She1 reduces dynein activity in cells^12,13^, which if compartmentally specified (*i.e*., to the mother cell), could have the capacity to polarize dynein-mediated spindle movements (*i.e*., to the daughter cell). Specifically, cells lacking She1 exhibit faster and more frequent dynein-mediated spindle movements, which are apparent from spindle movements coincident with astral microtubules sliding along the cell cortex^11-13,41^ (see Fig. 1G). Given our observations that She1 does not assist dynein-mediated nuclear migration into the bud neck (see Fig. 3D), but does impact the success rate for the spindle crossing the neck (see Fig. 1H), we sought to clarify this contradiction, and determine the basis by which She1 polarizes dynein activity by performing a more detailed assessment of dynein-mediated spindle movements. HU-arrested cells were imaged as described above, and dynein-mediated spindle movements were manually curated from automated spindle tracking data (using custom written MATLAB code; see Methods). We quantified various parameters of spindle movements from these data, including velocity, displacement per event, and relative degree of activity (as determined from the number of events per minute, and the total dynein-mediated spindle displacement, cell^-1^ min^-1^). Consistent with our previous *in vivo* and *in vitro* data, this analysis revealed that loss of She1 leads to increases in spindle velocity, displacement, and overall dynein activity (Fig. 4A – E).

**Figure 4.**
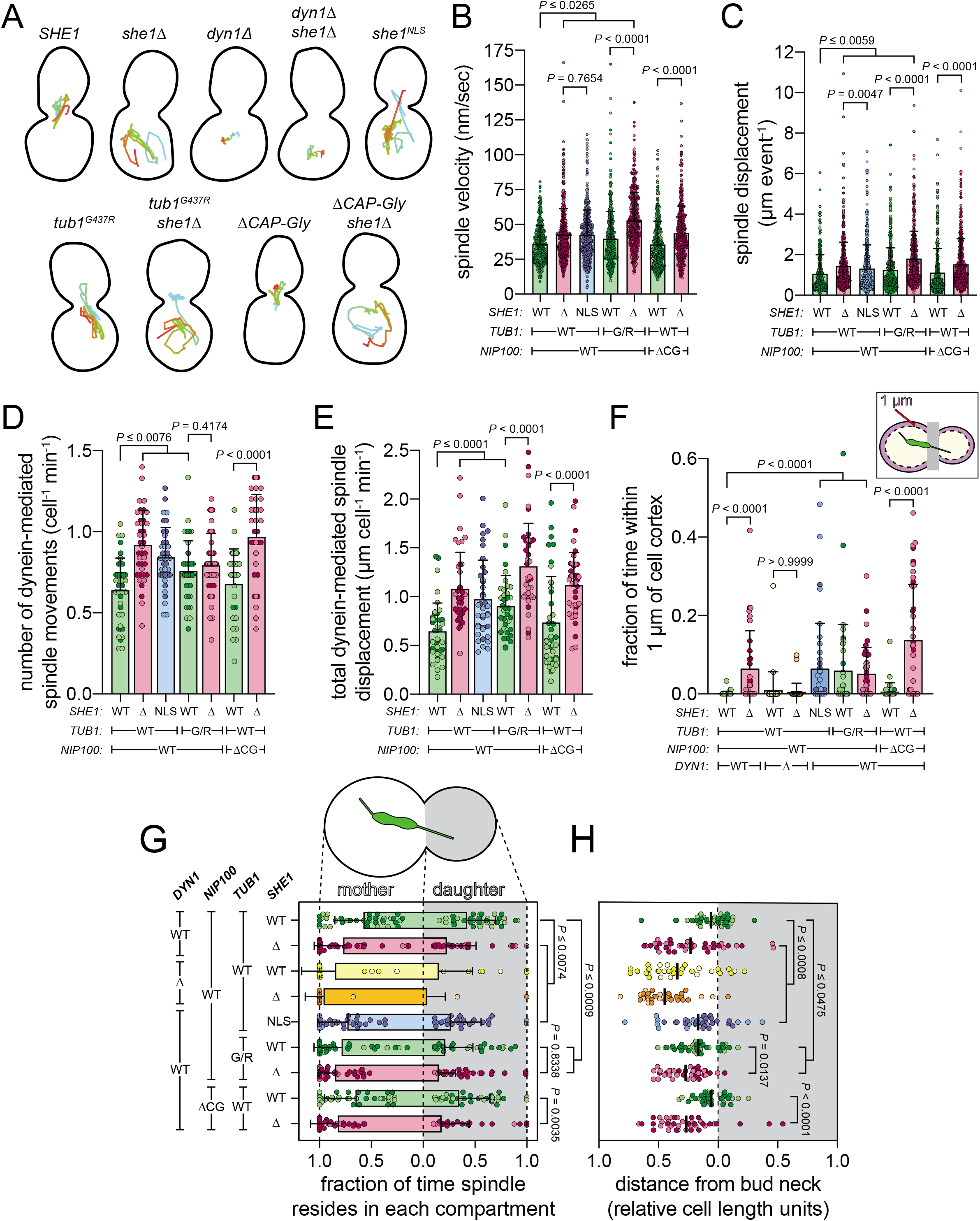
Astral microtubule-bound She1 promotes bud neck proximity of the spindle and reduces the quality and quantity of dynein-mediated spindle migration events. (A) Representative spindle tracks from indicated yeast strains. Hydroxyurea-arrested *kar9*Δ cells (along with indicated genotype) expressing GFP-Tub1 were imaged using confocal fluorescence microscopy, and a custom written MATLAB script was used to track the centroid of the spindle over time (time progression indicated by color gradient from cyan to red). (B – E) Dynein-mediated spindle movements were manually selected from the tracking data, from which velocity (B), displacement (C, per event; or, E, per minute), and the number of dynein-mediated spindle movements per minute (D) were obtained (mean ± standard deviation is overlaid with scatter plot of all data points; from left to right). For panels B – E, from left to right, n = 315 (40), 448 (40), 418 (38), 440 (40), 440 (40), 391 (40), and 440 (40) spindle movement events (number of cells) from 2 independent replicates. P values were calculated using the Mann-Whitney test. (F – H) By implementing two additional MATLAB scripts that determined the absolute position of the spindle centroid with respect to the cell cortex and bud neck (the latter of which also accounted for mother versus daughter cell information), we assessed: the relative number of spindle coordinates that reside within 1 μm of the cell cortex (F; mean ± standard deviation is overlaid with scatter plot from individual cells); the relative number of coordinates within the mother and daughter cell (G; circles represent fraction of time spindle resides within each compartment for individual cells); and, the mean position of the spindle centroid (along the longitudinal mother-daughter axis only) for each cell (H; circles represent mean spindle position for individual cells over the course of a 15 minute movie, and lines indicate mean values of all cells). For panel H, absolute positions were converted to relative values, where both mother and daughter cell were defined as 1 relative cell length unit (to account for differences in cell lengths). For panels F – H, from left to right (or top to bottom for G and H) n = 40, 40, 35, 35, 38, 40, 40, 40, and 40 cells from 2 independent replicates. P values were calculated using either the Mann-Whitney or an unpaired two-tailed Welch’s t-test (see Methods). For all panels, light and dark color hues indicate data points from independent replicates (“G/R”, *tub1^G437R^;* “ΔCG”, *nip100^ΔCAP-gly^*).

To gain additional insight into She1’s role in spindle positioning, we employed a custom written MATLAB code that identifies the relative position of the mitotic spindle over time with respect to the boundaries of budded yeast cells (*i.e*., mother versus daughter cell, and with respect to the bud neck or the cell cortex; to account for differences in cell lengths, the lengths of all mother and daughter cell were set to 1). This revealed that loss of She1 leads to: (1) the spindle spending a greater fraction of time within the mother cell, (2) an increased fraction of time during which the spindle resides within close proximity (≤ 1 μm) of the cell cortex, and (3) the spindle residing more distal from the bud neck (Fig. 4F – H). For example, whereas the spindle exhibits only a partial bias toward wild-type mother cells (57.6% in the mother, 42.4% in the daughter), with a mean position within very close proximity of the bud neck (a mean distance of 0.06 relative cell length units from the bud neck), the spindle spent a significantly larger fraction of time within *she1*Δ mother cells (77.1% in the mother, 22.9% in the daughter), and more distal from the neck (mean distance of 0.23 relative cell length units from the bud neck; Fig 4G and H).

Unlike the analysis described above (*i.e*., in Fig. 4A through E), which focused exclusively on dynein-dependent events, our spindle tracking data (in Figure 4F – H) include all positions in which the spindle resides over the course of a given movie. Therefore, to determine to what extent these spindle positions (*e.g*., within the daughter cell, or within close proximity of the bud neck) are due to dynein-dependent spindle migration, we repeated our analysis on cells deleted for the dynein heavy chain (*dyn1Δ*). As expected, these cells exhibited a large reduction (by 42.8% compared to *DYN1* cells; *P* < 0.0001) in spindle movements that were only minimally increased upon further deletion of She1 (by 16.5% compared to *dyn1*Δ; *P* = 0.0205; Fig. S3A). These data revealed that dynein and She1 are both required to promote spindle translocation in to the daughter cell, as spindles in both *dyn1*Δ and *dyn1*Δ *she1*Δ cells spent a very low fraction of time within the daughter cell (15.1% and 3.8% for *dyn1*Δ and *dyn1*Δ *she1*Δ, respectively; Fig. 4G). We also noted a very low fraction of time the spindle resided within close proximity of the cell cortex in these mutants, indicating this phenomenon is indeed dynein-dependent (Fig. 4F). Finally, although deletion of dynein leads to an increased distance of the spindle from the bud neck (0.34 relative cell length units), this value was slightly increased upon deletion of She1 (to 0.45 relative cell length units; *P* = 0.0647), suggesting that She1 may play a dynein-independent role in promoting spindle-bud neck proximity (Fig. 4H). Taken together, these data indicate that She1 attenuates the initiation of dynein-mediated spindle movements, and ensures that the mitotic spindle remains distal from the cell cortex, and within close proximity of the bud neck. Given the biased residence of the spindle within *she1*Δ mother cells (Fig. 4G and H), these data suggest that She1 may specifically attenuate dynein activity within the mother cell by an unknown mechanism.

### She1 affects spindle position in a manner that requires its binding to cytoplasmic microtubules

Our data indicate that She1 affects at least two distinct aspects of cellular dynein activity: (1) the quality of dynein motility (*e.g*., it reduces the velocity and displacement of dynein-mediated spindle movements; Fig. 4B and C), and (2) initiation of dynein motility (*e.g*., it reduces the number and extent of dynein-mediated events; Fig. 4D and E). Our previous reconstitution experiments revealed that She1 reduces dynein velocity via simultaneous interactions between the dynein microtubule-binding domain and the microtubule^10,13^, which may account for the spindle velocity reduction phenotype in cells, and may potentially also underly the increased stall times we observed in our optical trapping experiments (see Fig. 2E). However, none of our *in vitro* data revealed a She1-mediated reduction in microtubule binding by dynein, raising the question of how She1 may inhibit initiation of dynein-mediated spindle translocation events. To gain insight into the mechanism underlying this phenomenon, and to improve our understanding of the velocity reduction effect, we employed our detailed spindle dynamics assays in combination with a series of various mutants.

In addition to binding to astral microtubules in the cytoplasm^11^, She1 also localizes prominently to spindle microtubules in the nucleus^42^, where it has been proposed to play several roles (*e.g*., in promoting spindle disassembly^37^, maintaining spindle microtubule stability^43^, and potentially in affecting kinetochore function^42^). Given the role of She1 in these spindle functions, it is conceivable that loss of She1 impacts spindle movements from within the nucleus. For instance, since loss of She1 can impact spindle stability, it is conceivable that dynein-mediated outward forces that originate from the cell cortex could be impacted by the loss of inward-directed forces provided by spindle stabilizing, microtubule-binding factors (*e.g*., She1). To determine whether it is the nuclear or cytoplasmic pool of She1 that is responsible for affecting the various spindle motility parameters, we added a strong nuclear localization signal (from the SV40 large T antigen) to the C-terminus of endogenous She1 (at the genomic locus). Immunoblotting confirmed that She1^NLS^ was expressed at levels similar to that of wildtype She1 in cells (Fig. S2A). Using purified proteins and fluorescence microscopy-based binding assays (Fig. S2B and C), we also found that She1^NLS^ binds to yeast microtubules (assembled from tubulin purified from yeast) with very similar affinity as wild-type She1 (Fig. S2D and E; K_D_ = 854 nM and 806 nM for wild-type and NLS, respectively).

Analysis of spindle movements in *she1^NLS^* cells revealed a phenotypic signature that was almost identical to *she1*Δ cells. For instance, the velocity and run length values for spindle movements were increased to a similar extent in *she1*Δ and *she1^NLS^* cells (Fig. 4A – C). We also noted a similar increase in the relative extent of dynein activity (Fig. 4D and E), and spindle positioning metrics that closely mirrored *she1*Δ cells (*e.g*., increased residence of the spindle in the mother cell, in close proximity to the cell cortex, and distal from the bud neck; Fig. 4F – H). These data indicate that She1 affects both the quality and quantity of dynein-mediated spindle movements from within the cytoplasm.

As noted above, our previous work demonstrated that She1 must be bound to microtubules to reduce dynein velocity *in vitro^10^*. To determine whether this is the case in cells, and to assess the contribution of microtubule-bound She1 to the initiation of dynein-mediated spindle movements, we employed cells expressing a mutant a-tubulin (Tub1^G437R^, with a Gly to Arg substitution immediately adjacent to the C-terminal unstructured E-hook; referred to as “G/R” in Fig. 4) to which She1 binds to a significantly reduced extent (55.7% reduction in apparent microtubule-binding in *tub1^G437R^* cells^44^). Consistent with the notion that She1 must bind microtubules to affect dynein motility, *tub1^G437R^* mutant cells exhibited increased spindle velocity and displacement values that roughly scale with the relative reduction in She1-microtubule binding in these cells (Fig. 4B and C). Moreover, *tub1^G437R^* cells exhibit an increase in the relative extent of dynein activity (Fig. 4A, D and E), a mean spindle position distal from the bud neck (Fig. 4H), and increased fractions of time during which the spindle resides in close proximity of the cell cortex and within the mother cell (Fig. 4F and G). Further deletion of She1 in *tub1^G437R^* cells led to more robust phenotypes, consistent with the partial ability of She1 to bind to microtubules in the mutant cells. Thus, astral microtubule-bound She1 indeed reduces the quality and quantity of dynein-mediated spindle movements. Given the increased residence of the spindle in the mother cell in both *tub1^G437R^* and *she1^NLS^* cells, these data indicate that astral microtubule-bound She1 reduces initiation of dynein activity specifically in the mother cell.

### She1 does not preclude initiation of dynein-mediated spindle movements via the dynactin CAP-gly microtubule-binding domain

In addition to directly impacting dynein motility (*in vitro* and *in vivo*), She1 has also been implicated in affecting the interaction between dynein and dynactin, a multisubunit complex that is required for dynein activity in cells^11,12,24^. Although the mechanism by which She1 performs this activity and the impact of this regulation on cellular dynein function is unclear, we wondered whether some of our observed effects on spindle migration are due to She1 affecting the dynein-dynactin interaction. Previous studies have found that human dynactin promotes a motility-competent configuration of dynein, and also stimulates microtubule-binding of the dynein-dynactin complex^45,46^. The manner by which dynactin enhances microtubule binding of the human dynein-dynactin complex is due in large part to the N-terminal <u>C</u>ytoskeleton-<u>A</u>ssociated Protein-<u>gly</u>cine rich (CAP-gly) microtubule-binding domain on the dynactin subunit p150^Glued^ (Nip100 in budding yeast)^45^. Thus, it is feasible that enhanced dynein-dynactin binding could lead to an increase in the initiation of dynein-mediated spindle movements. Given that deletion of any of the dynactin subunits in budding yeast severely compromises dynein activity^17^, we chose to employ a Nip100^ΔCAP-gly^ mutant (“ΔCG”) that has been shown to have only minor effects on dynein activity^39^.

With the exception of a small reduction in the time the spindle spends within the daughter cell (42.4% for wild-type versus 34.8% for *nip100^ΔCAP-gly^* cells; Fig. 4G), our analysis of spindle movements in *nip100^ΔCAP-gly^* mutant cells revealed no significant impact on dynein function (Fig. 4A – H). These data contrast with prior observations in budding yeast^39^, and instead suggest that the Nip100 CAP-gly domain plays a very minor role in dynein-mediated spindle movements. Moreover, additional deletion of She1 (*i.e*., *she1Δ nip100^ΔCAP-gly^* cells) led to a phenotypic signature largely indistinguishable from *NIP100 she1*Δ cells. Thus, although we cannot rule out the possibility that She1 affects cellular dynein activity in a dynactin-dependent manner, these data indicate that the dynactin microtubule-binding domain is not involved in this mode of regulation.

### She1-mediated attenuation of dynein activity requires an interaction with the dynein microtubule binding domain

We previously found that She1 affects *in vitro* dynein motility through simultaneous interactions with the microtubule and the dynein microtubule-binding domain (MTBD)^10^. We thus sought to determine the importance of the She1-dynein MTBD interaction on the various aspects of spindle migration. To this end, we employed a chimeric dynein mutant (Dyn1^mMTBD^; in which the yeast dynein MTBD was substituted for the corresponding mouse sequence) that exhibits reduced affinity for She1, and reduced sensitivity to She1 in single molecule assays^10^. Given the *dyn1^mMTBD^* allele possesses a C-terminal 3YFP tag (in contrast to the untagged *DYN1* allele employed in the above assays), we compared spindle dynamics of this mutant to cells expressing wild-type Dyn1-3YFP. Although the *DYN1-3YFP* allele supports normal spindle positioning^23^, we noted that these cells exhibited somewhat reduced spindle velocity values (35.9 nm/sec for *DYN1* versus 29.9 nm/sec for *DYN1-3YFP; P* < 0.0001; Fig. S3B). Moreover, although deletion of She1 led to similar alterations in most spindle migration parameters in *DYN1-3YFP* cells (compared to *DYN1* cells), many of the *she1*Δ-mediated changes were somewhat attenuated (*e.g*., velocity, displacement and activity increases, and mother/daughter residence; Fig. S3C – G), suggesting that Dyn1-3YFP – which may have compromised motility metrics compared to Dyn1 – is less susceptible to She1-mediated inhibition.

Unexpectedly, we noted a *she1*Δ-mediated increase in the quality of dynein-mediated spindle motility in *dyn1^mMTBD^-3YFP* cells (velocity, run length and cortical proximity; Fig. 5A – C, and F), suggesting that in-cell modulation of dynein motility by She1 does not occur through binding to the dynein MTBD. However, the *she1*Δ-dependent increase in the extent of dynein activity in *dyn1^mMTBD^* cells was significantly attenuated compared to wild-type cells, indicating that She1’s inhibition of dynein activity initiation does indeed occur in a manner that is dependent on the interaction between She1 and the dynein MTBD (Fig. 5D and E). These data contrast with our *in vitro* findings, which indicate that this interaction is required for the velocity reduction effect with purified dynein^10^, and suggest that cortical Num1-dynein-dynactin complexes are affected differently by She1 than dynein alone. They also indicate that the increased cortical proximity of the spindle is largely a consequence of altered motility quality in *she1*Δ cells (*i.e*., velocity and run length), and is not due to an enhanced initiation of motility.

**Figure 5.**
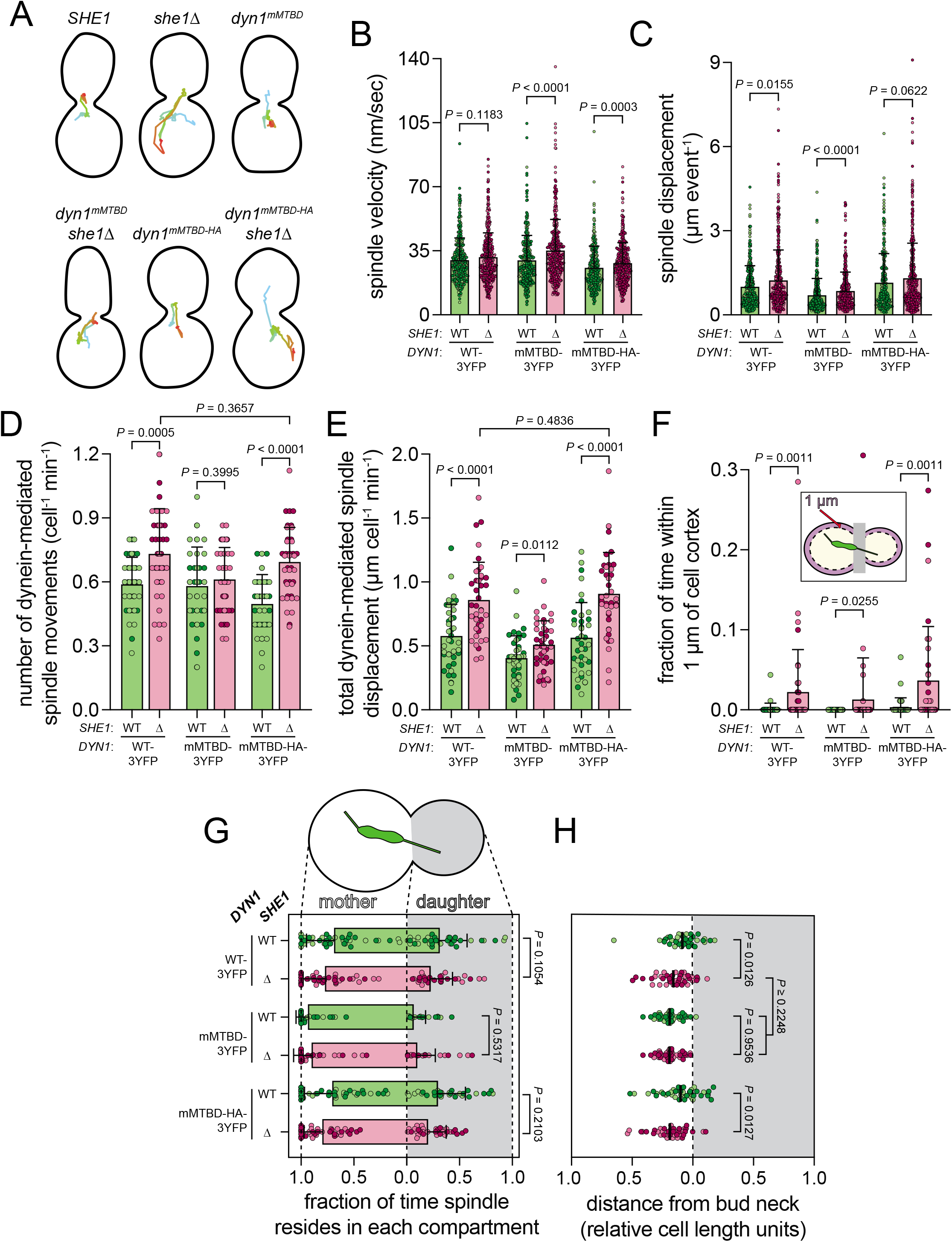
She1 maintains bud neck proximity of the spindle and attenuates initiation of dynein activity through interactions with the dynein MTBD. (A) Representative spindle tracks from indicated yeast strains. Hydroxyurea-arrested *kar9*Δ cells (along with indicated genotype) expressing GFP-Tub1 were imaged using confocal fluorescence microscopy, and a custom written MATLAB script was used to track the centroid of the spindle over time (time progression indicated by color gradient from cyan to red). (B – E) Dynein-mediated spindle movements were manually selected from the tracking data, from which velocity (B), displacement (C, per event; or, E, per minute), and the number of dynein-mediated spindle movements per minute (D) were obtained (mean ± standard deviation is overlaid with scatter plot of all data points; from left to right). For panels B – E, from left to right, n = 346 (40), 418 (40), 351 (40), 361 (40), 265 (40), and 417 (40) spindle movement events (number of cells) from 2 independent replicates. P values were calculated using the Mann-Whitney test. (F – H) Plots depicting the relative number of spindle coordinates that reside within 1 μm of the cell cortex (F; mean ± standard deviation is overlaid with scatter plot from individual cells), the relative number of coordinates within the mother and daughter cell (G; circles represent fraction of time spindle resides within each compartment for individual cells), and, the mean position of the spindle centroid (along the longitudinal mother-daughter axis only) for each cell (H; circles represent mean spindle position for individual cells over the course of a 15 minute movie, and lines indicate mean values for all cells). See Figure 4 for additional information. For panels F – H, from left to right (or top to bottom for G and H) n = 40, 40, 40, 40, 40, and 40 cells from 2 independent replicates. P values were calculated using either the Mann-Whitney or an unpaired two-tailed Welch’s t-test (see Methods). For all panels, light and dark color hues indicate data points from independent replicates (“WT-3YFP”, *DYN1-3YFP;* “mMTBD-3YFP”, *dyn1^mMBTD^-3YFP*; “mMTBD-HA-3YFP”, *dyn1^mMTBD-HA^-3YFP*).

Our prior *in vitro* work revealed that She1 exhibits higher affinity for the nucleotide-free conformational state of the dynein motor domain (the apo state)^10^. Given She1’s interaction with the dynein MTBD, and its preferential binding to the high microtubule affinity state of the MTBD^10^, this indicates that She1 exhibits higher affinity for dynein when it is bound to microtubules during a processive run^47,48^. Although the purpose of this binding specificity is not understood, it may play a role in the mechanism by which She1 regulates dynein activity. We previously found that Dyn1^mMTBD^ exhibits a lower affinity for microtubules than the wild-type yeast motor^10^, suggesting that the basis for the partial insensitivity of the chimera to She1 may be a consequence of conformational differences between the two MTBDs (*i.e*., mouse versus yeast). To determine whether this is the case, we introduced two charge-reversal point mutations into Dyn1^mMTBD^ (E3289K and E3378K; in the coiled-coil and MTBD, respectively), which have been previously reported to increase dynein-microtubule binding affinity^49^. This mutant would be expected to spend a greater fraction of time in a microtubule-bound conformational state – to which She1 would be predicted to preferentially bind – and would potentially restore She1 sensitivity to the chimeric mutant. Cells expressing this mutant (Dyn1^mMTBD-HA^, for <u>h</u>igh <u>a</u>ffinity) exhibited similar activity metrics to wild-type cells (Fig. 5D and E), but moved their spindles significantly slower than both *DYN1* and *dyn1^mMTBD^* cells (Fig. 5B; *P* < 0.0001), as would be expected for a motor that spends more time in a microtubule-bound state. Interestingly, the *she1*Δ-mediated increase in the initiation of dynein-mediated spindle movement events was completely restored in *dyn1^mMTBD^-^HA^ she1*Δ cells (compared to *DYN1 she1*Δ cells; *P* ≥ 0.3657 for sliding events and spindle displacement; Fig. 5D and E), indicating that the She1-mediated inhibition of dynein activity indeed requires a microtubule-bound conformational state of the dynein MTBD.

Analysis of tracking data revealed an increased fraction of time during which the spindle resides within the mother cell, and a mean position more distal from the bud neck as a consequence of She1 deletion in *DYN1* and *dyn1^mMTBD-HA^*, but not *dyn1^mMTBD^* cells (Fig. 5G and H). In light of the varying extents to which loss of She1 affects the dynein activity parameters for each of the mutants, these data suggest that She1-mediated inhibition of the initiation of dynein-mediated spindle movements (as shown in Fig. 5D and E), and not its effect on the quality of movement (as shown in Fig. 5B and C), is the key determinant that dictates She1’s ability to promote bud neck proximity, and daughter cell-directed spindle movements.

### She1 inhibits the initiation of dynein-mediated spindle movements in the mother cell

We next sought to determine whether She1 directly affects the initiation of dynein-mediated spindle movements in a spatially confined, compartment-specific manner (*i.e*., mother versus daughter cell), which could potentially account for the mother cell bias in spindle residence. Astral microtubule plus ends make contacts with the cell cortex as they dynamically sample the mother and daughter cell compartments, with a subset of these ‘cortical contacts’ resulting in a dynein-mediated spindle movement (referred to as a ‘productive event’; Fig. 6A). Initiation of a productive event is thought to occur coincident with offloading and activation of the dynein-dynactin complex at Num1 sites^24,25^. We wondered whether the increased frequency of dynein-mediated spindle movement events in *she1*Δ cells is a consequence of (1) more astral microtubule contacts with the cell cortex (*e.g*., due to changes in microtubule dynamics), (2) or more such contacts transitioning into a productive event. To address these questions, we separately counted the total number of cortical contacts and productive events in mother and daughter cells from live movies.

**Figure 6.**
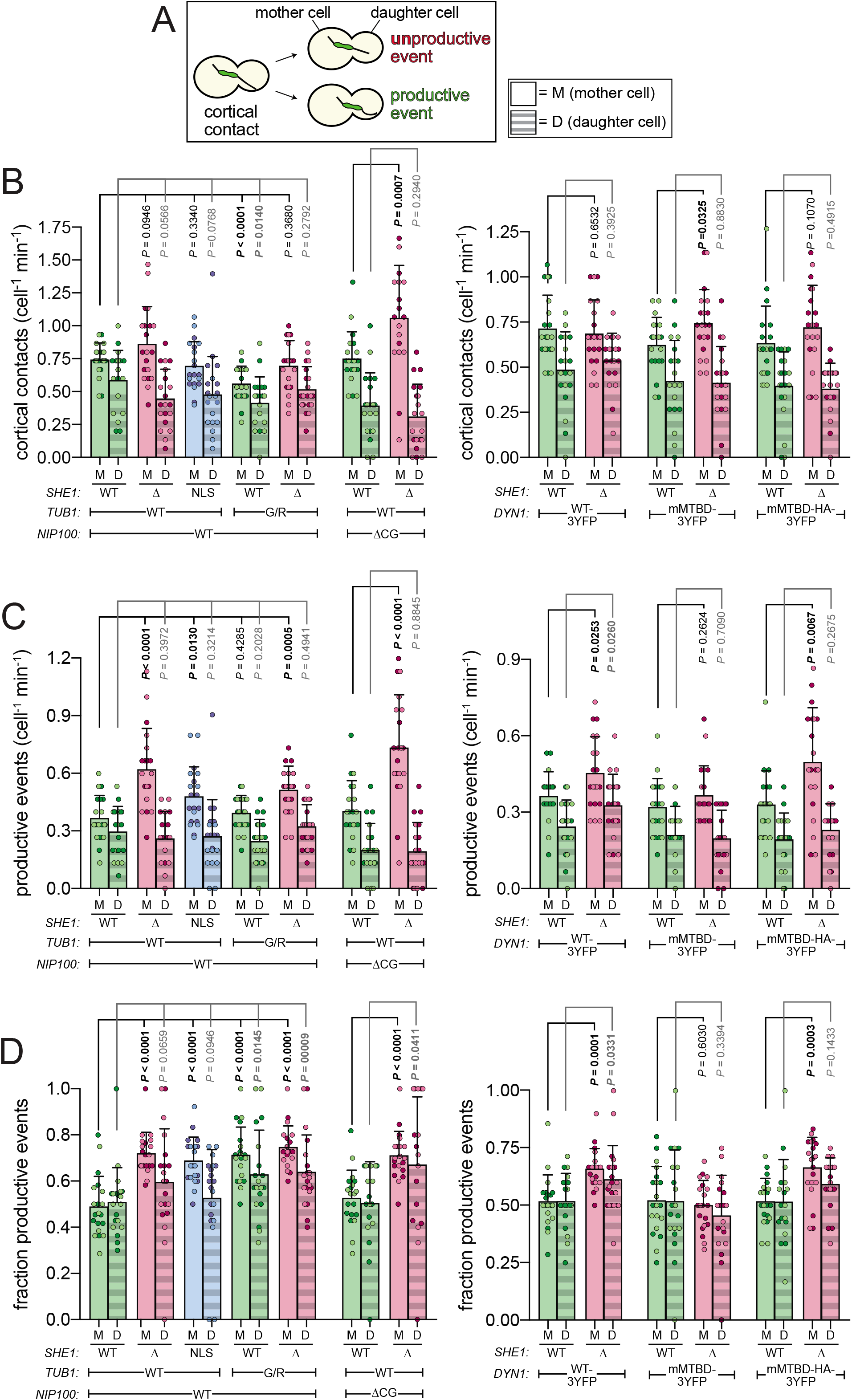
She1 attenuates initiation of dynein activity in the mother cell through interactions with microtubules and the dynein MTBD. (A) Cartoon schematic depicting events quantitated in panels B – D. (B – D) Hydroxyurea-arrested *kar9*Δ cells (along with indicated genotype) expressing GFP-Tub1 were imaged using confocal fluorescence microscopy, and both XY and XZ maximum intensity projections were used to quantitate the frequency with which astral microtubule plus ends contact the cell cortex (B), the frequency of ‘productive’ dynein-mediated spindle translocation events (C), and the fraction of cortical contacts that convert to a productive event (D; for all strains, n = 10 cells from 2 independent replicates). P values were calculated using either the Mann-Whitney or an unpaired two-tailed Welch’s t-test (see Methods). For all panels, light and dark color hues indicate data points from independent replicates (“G/R”, *tub1^G437R^;* “ΔCG”, *nip100^ΔCAP-gly^;* “WT-3YFP”, *DYN1-3YFP;* “mMTBD-3YFP”, *dyn1^mMBTD^-3YFP*; “mMTBD-HA-3YFP”, *dyn1^mMTBD-HA^-3YFP*).

This analysis revealed that the number of cortical contacts was significantly greater in daughter than mother cells (Fig. 6B), consistent with previous observations that the daughter-oriented SPB nucleates more and longer microtubules^50–53^. Deletion of She1 had very little impact on the total number of cortical contacts in either mother or daughter cells (Fig. 6B). This was largely the case for most of the mutants, with the exception of *nip100^ΔCAP-gly^ she1*Δ cells, which exhibited significantly more contacts in the mother cell than the parent *nip100^ΔCAP-gly^* strain (*P* = 0.0007). In spite of the unchanged number of microtubule-cortex encounters in *she1*Δ cells, the number of dynein-mediated sliding events was significantly greater in *she1*Δ mother, but not *she1*Δ daughter cells with respect to wild-type cells (*P* < 0.0001 and *P* = 0.3972, respectively; Fig. 6C). Moreover, plotting the relative fraction of cortical contacts that transition to a productive events in each compartment revealed that She1 indeed dampens the initiation of a dynein-mediated spindle migration events specifically in the mother cell (Fig. 6D). Specifically, whereas an approximately equal fraction of cortical contacts in wild-type mother and daughter cells result in a productive event (49.0% and 51.0% in mother and daughter, respectively), these values were increased to 72.0% in *she1*Δ mother cells (*P* < 0.0001), but only 59.6% in *she1*Δ daughter cells (*P* = 0.0659). Notably, we observed a very similar pattern of disproportionate increases in the fraction of productive events in mother cells in all those mutants that exhibited a biased enrichment of spindle residencies in the mother cell, including *she1^NLS^, tub1^G437R^, tub1^G437R^ she1Δ*, and *dyn1^mMTBD-HA^ she1*Δ cells. The only exceptions were *dyn1^mMTBD^* cells, which showed no *she1*Δ-dependent increase in dynein activity in mother or daughter cells (*P* = 0.6030 and 0.3394, respectively), and *nip100^ΔCAP-gly^ she1*Δ cells, which exhibited a broad distribution of values, and an overall increase in the mean activity in both mother and daughter cells upon She1 deletion (*P* < 0.0001 and *P* = 0.0411, respectively). Taken together, these data indicate that astral microtubule-bound She1 dampens the initiation of dynein-mediated spindle movement events specifically in the mother cell in a manner that requires an interaction with the microtubule-bound conformation of the dynein MTBD. These findings likely account for the biased spindle residence in mother cells upon loss of She1.

### She1 localizes preferentially to astral microtubules within the mother cell

Given the importance of astral microtubule-She1 binding in preventing the initiation of dynein-mediated spindle movements in the mother cell, we wondered whether She1 preferentially localizes to astral microtubules in this compartment. To circumvent the difficulty in visualizing endogenous levels of She1-3GFP (likely due to low cellular concentrations), we overexpressed She1-3GFP using the galactose-inducible promoter, GAL1p. To be able to correlate She1 localization patterns to our spindle dynamics data, we arrested *GAL1p:SHE1-3GFP* cells in HU, as described above. Live cell imaging revealed a striking difference in the localization pattern of She1 to microtubules between mother and daughter cells. Specifically, we observed that 82.5% of mother cells had apparent She1-3GFP along astral microtubules, compared to only 20.0% of daughter cells (note that only those cells with astral microtubules apparent in both mother and daughter cells were selected for quantification; Fig. 7A and B). We confirmed the preferential localization of She1 to astral microtubules in the mother cell by comparing the fluorescence intensity values of She1-3GFP on astral microtubules in each compartment, which revealed a 3-fold greater degree of microtubule-binding in mother cells compared to daughter cells (Fig 7C). We also ensured this was not a consequence of differences in microtubule polymer mass (*e.g*., due to microtubule bundling) by comparing the relative intensity of She1-3GFP to that of mRuby2-Tub1 on individual microtubules, which revealed a similar 3.7-fold greater binding to mother cell microtubules (Fig. 7D). These data indicate that She1 exhibits preferential binding to microtubules in the mother cell, which likely accounts for the biased inhibition of dynein-mediated spindle movements in this compartment.

**Figure 7.**
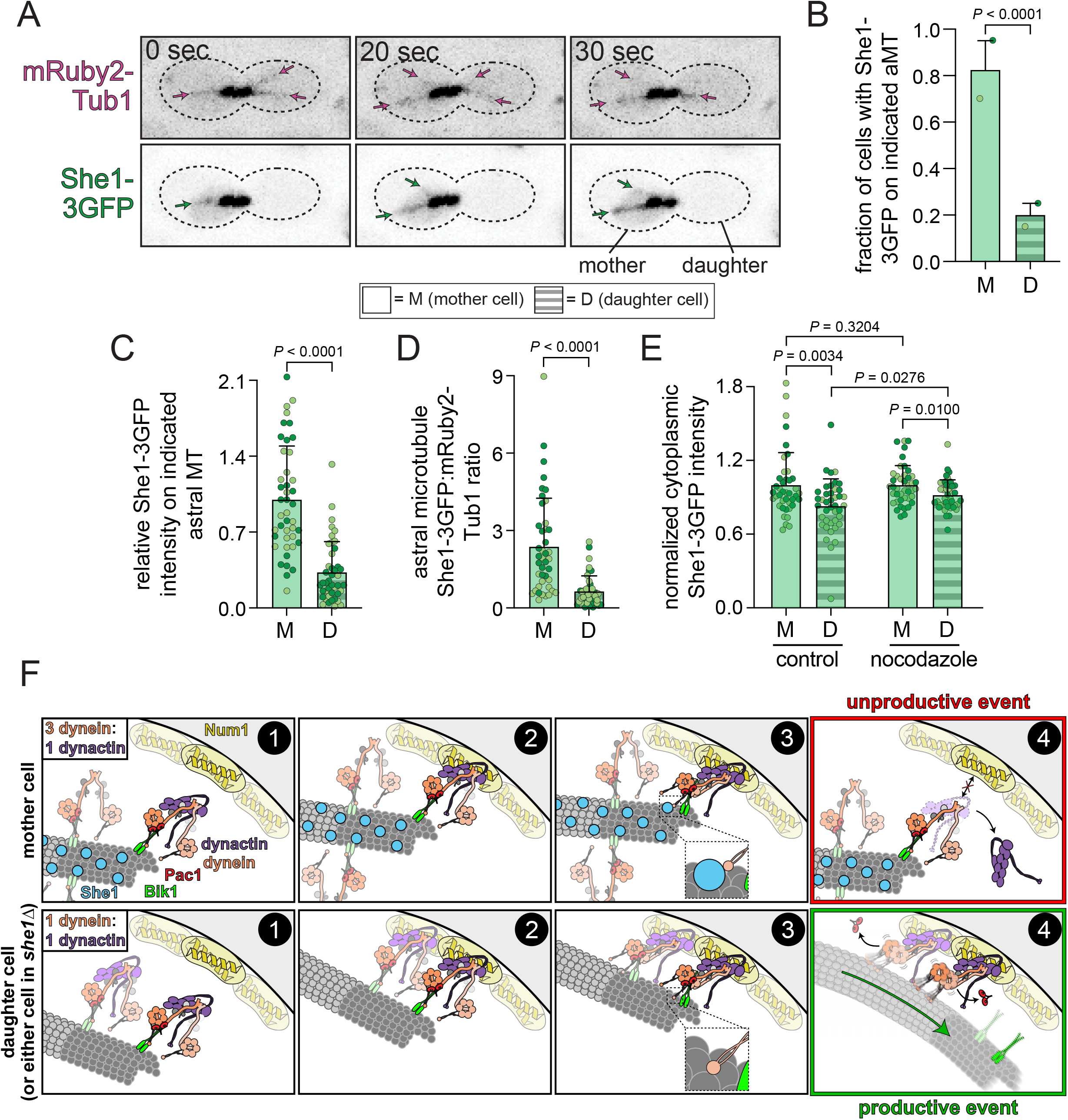
She1 preferentially localizes to astral microtubules in the mother cell. (A) Representative time-lapse confocal fluorescence microscopy images of a hydroxyurea-arrested cell expressing mRuby2-Tub1 and overexpressing She1-3GFP. Note the presence of She1-3GFP on astral microtubules within the mother cell (bottom; green arrows), but not the daughter cell. (B) Plot depicting the fraction of cells with apparent She1-3GFP on either mother or daughter astral microtubules. Only cells with apparent astral microtubules in both mother and daughter cells were assessed. Circles represent fractions from independent replicates, and bars represent mean ± standard deviation (n = 40 cells from 2 independent replicates). P value was calculated from Z-score as described previously^76^. (C) Plot depicting background-corrected She1-3GFP intensity values on astral microtubules in mother and daughter cells. Astral microtubules were identified and delineated from mRuby2-Tub1 fluorescence, thus permitting us to measure She1-3GFP intensity in cells with little to no apparent fluorescence. (D) Plot depicting She1-3GFP intensity relative to mRuby2-Tub1 intensity. Fluorescence intensities of each were measured, and respective values from individual microtubules were divided to obtain relative values (She1-3GFP divided by mRuby2-Tub1). For panels C and D, n = 43 cells from 2 independent replicates. As with panel B, only cells with apparent astral microtubules in both mother and daughter cells were used for intensity measurements. (E) Plot depicting background-corrected cytoplasmic fluorescence intensity values for She1-3GFP from either control cells, or those treated with 100 μM nocodazole for 30 minutes (which completely depolymerizes astral microtubules; see Fig. S4; n = 42 and 43 control and nocodazole-treated cells, respectively, from 2 independent replicates). For panels C – E, circles represent values from individual microtubule measurements, and bars are mean ± standard deviation. P values were calculated using either the Mann-Whitney or an unpaired two-tailed Welch’s t-test (see Methods). For panels B – E, light and dark color hues indicate data points from independent replicates. (F) Model for She1-mediated attenuation of dynein-mediated spindle migration activity. Plus end-associated dynein (step 1) – 1 of 3 of which are associated with dynactin in the presence of She1 (top), but all of which are associated with dynactin in its absence (bottom)^12^ – contacts cortical Num1 (step 2), which promotes dynein-microtubule binding (step 3; potentially by reorienting the motor domains into a parallel, motility-competent configuration^46^). In the absence of She1 (bottom), dynein dissociates from the plus end-targeting machinery (*i.e*., via release of Pac1)^25^, and is thus activate to translocate the spindle (step 4, bottom). However, upon She1-dynein MTBD binding (top), the dynein-dynactin complex dissociates, which leads to Num1-dissociation, terminating the offloading event (step 4, top).

We hypothesized that the biased localization of She1 to astral microtubules within the mother cell is a consequence of either asymmetric enrichment within the cytoplasm of the mother cell, or preferential microtubule-binding activity in this compartment. To determine whether She1 is enriched in the mother cell cytoplasm, we measured the fluorescence intensity values for regions within the cytoplasm (*i.e*., regions not including microtubules) of untreated cells, and those treated with the microtubule depolymerizing agent nocodazole (see Fig. S4). This analysis revealed a small enrichment of She1-3GFP in the untreated mother cells (18% greater intensity in mother cells; Fig. 7E) that was somewhat reduced in nocodazole-treated cells (8% enrichment in mother cells). Given the large difference in microtubule-binding between the two compartments (3 to 3.7-fold) compared to these small differences in cytoplasmic intensities, these data suggest that the biased localization of She1 to mother cell-associated astral microtubules is largely achieved through differential microtubule binding activity in the two compartments, possibly through the modulation of She1 affinity for microtubules.

## DISCUSSION

Here, we have revealed the means by which She1 affects dynein activity in cells, and how it polarizes dynein-mediated spindle movements toward the daughter cell. Specifically, we find that She1 precludes the initiation of dynein-mediated spindle movements, which is apparent from a reduction in the conversion of astral microtubule-cortex encounters to productive spindle translocation events. This mode of inhibition, which occurs more prominently in mother cells, requires She1 binding to astral microtubules and the dynein MTBD. The compartment-specific inhibition mechanism is likely achieved as a result of She1’s preferential binding to astral microtubules within mother cells with respect to daughter cells. Although our prior *in vitro* work revealed the She1-dynein MTBD interaction, and demonstrated the importance of this interaction for velocity reduction in single molecule assays with purified dynein alone^10^, the nature of in-cell inhibition of dynein activity appears to occur through an entirely distinct mechanism that is unrelated to She1’s ability to reduce the velocity of dynein, or reduce its microtubule-dissociation rate *in vitro*. Thus, although She1 can promote *in vitro* dynein-microtubule binding in the absence^10^ and presence of load (see Fig. 2), it appears that She1 in fact precludes productive dynein-microtubule encounters in cells. Given that dynein functions in concert with dynactin and Num1 at the cell cortex, our data indicate that She1 affects the Num1-dynein-dynactin complex in a manner that is distinct from dynein alone.

Our new data have forced us to revise the model by which She1 affects cellular dynein activity. Although She1 interactions with the dynein MTBD and microtubules are required for the She1-mediated reduction in the initiation of dynein-mediated spindle movements in cells, neither interaction appears to be required for the velocity reduction effect (*i.e*., see *dyn1^mMTBD^* and *tub1^G437R^* data; Fig. 4 and 5). How might She1 reduce the quality of dynein-mediated spindle movements (*e.g*., velocity and displacement event^-1^)? She1 deletion leads to more intact dynein-dynactin complexes at plus ends^11,12^ (from ~3 dynein:1 dynactin per plus end, to ~1:1 in *she1*Δ cells^12^; see Fig. 7F, top and bottom). This mechanism, which may occur in a manner that is independent of the She1-dynein MTBD interaction, would result in the offloading of a greater number of active motor complexes per astral microtubule-cortex contact (see Fig. 7F, bottom). As a result, this would provide more active motors for a given spindle pulling event, and thus greater velocities and displacements. Notably, data from *dyn1^mMTBD^* cells – in which the She1-mediated reduction in the initiation of events is uncoupled from its ability to reduce spindle velocity – indicate that the former is the activity required to polarize dynein activity. Thus, while the cortical proximity phenotype appears to be a consequence of She1 reducing the velocity and/or displacement of each dynein-mediated spindle movement, the She1-mediated spindle positioning phenotypes (*i.e*., toward the daughter cell and within proximity of the bud neck) appear to be due to She1 attenuating the initiation of dynein activity in the mother cell.

How does She1 inhibit the initiation of dynein-mediated spindle movements? We find that She1 only inhibits initiation of dynein motility when it is in the microtubule-bound conformational state (compare *dyn1^mMTBD^* to *dyn1^mMTBD-HA^* cells; Fig. 5), for which She1 exhibits higher affinity^10^. Initiation of a dynein-mediated spindle movement event in budding yeast is likely immediately preceded by the offloading of dynein-dynactin complexes from microtubule plus ends to cortical Num1 receptor sites, from where they engage with astral microtubules to move the spindle^20,23,24,54^. Dynein does not contact the plus end directly (*i.e*., is MTBD-independent), but rather interacts indirectly with the plus end-associated factors Pac1 and Bik1 (homologs of human LIS1 and CLIP170; Fig. 7F, step 1)^25^. However, we recently proposed a model whereby dynein does make direct contact with the microtubule during the offloading process, and that this contact is required to disengage dynein from the plus end-targeting machinery (*i.e*., Bik1 and Pac1), thus permitting spindle translocation^25^ (see Fig. 7F; step 3). If true, this would be the first moment in the dynein activation process at which microtubule-bound She1 encounters the dynein MTBD. Thus, we posit the following model to account for She1-mediated inhibition of dynein activity: (1) plus end-bound dynein-dynactin complexes associate with cortical Num1, but do not yet dissociate from plus ends (Fig. 7F; step 2); (2) as a consequence of their association with Num1, dynein-dynactin switch to an active, motile state^25^ (which may be due to the dynein motor heads orienting in a parallel manner^46^), and the dynein MTBD makes direct contacts with the microtubule and thus microtubule-bound She1 (Fig. 7F, top; step 3); (3) although in the absence of She1, the dynein-microtubule binding step leads to dynein-dynactin dissociating from the plus end-binding proteins^25^ (thus permitting spindle migration activity; Fig. 7F, bottom; steps 3 and 4), the dynein MTBD-She1 interaction causes dynein to dissociate from dynactin, which breaks dynein’s contact with Num1 (due to the reliance of dynein on dynactin for Num1 binding^55^), thus terminating the offloading event (Fig. 7F, top; step 4). This model is supported by our prior observation that deletion of She1 leads to enhanced offloading of dynein to cortical sites^24^.

How might She1-dynein MTBD binding lead to dynein-dynactin dissociation? In light of the large distance between the dynein MTBD and the dynein-dynactin contact points (at the dynein intermediate chain-p150/Nip100 interaction surface^56–59^, and the dynein heavy chain N-terminus-Arp1 filament interaction surface^60,61^), She1 likely exerts allosteric effects to disrupt these contacts. One potential mechanism may employ the autoinhibited conformational state of dynein, which, when engaged, reduces dynein-dynactin affinity^46^. Thus, the She1-dynein MTBD contacts may directly promote and/or stabilize the autoinhibited conformation, which is mediated in part by contact points in the coiled-coil stalk within the heavy chain^46,54^, which is in close proximity to the MTBD.

In addition to its role in spindle positioning (see Fig. 1A and B), our cell cycle progression data also revealed a potential role for She1 in the spindle position checkpoint (SPC), as deletion of She1 leads to no checkpoint delay in cells with spindle positioning defects (see Fig. 1F). The SPC ensures that mitotic exit only occurs in cells in which the mitotic spindle is properly positioned (with one SPB each in mother and daughter cell, respectively; see Fig. 1A). This checkpoint functions by inhibiting the activity of the mitotic exit network (MEN), which promotes exit by a complex signaling network (reviewed in^27^). In addition to potentially functioning in the SPC, She1 shares two other noteworthy similarities with SPC effectors: prominent localization to the bud neck^42^, and an asymmetric localization pattern (see Fig. 7). For example, like She1, the Kin4 kinase and its activating kinase Elm1 – both critical SPC effectors – localize to the bud neck, where they promote SPC function^32–36^. Moreover, Kin4 and the guanine nucleotide exchange factor Lte1 each localize only to mother and daughter cell compartments, respectively, where they sense and respond to the position of the SPBs^28,31,33,34,62^. Although we find that microtubule-bound She1 is important to regulate dynein activity, the role of the bud neck-localized pool of She1 remains unknown, but may be involved in SPC function. Thus, She1 may be the first known molecule that functions in both directly promoting appropriate spindle position, and in conveying to the spindle position status to the cell cycle progression machinery.

The means by which asymmetric localization of MEN effectors and She1 are achieved is unclear (although one study suggests a septin-dependent diffusion barrier for membrane-associated Lte1^63^); however, it is possible that they involve similar mechanisms. In light of our data that She1 asymmetry is likely imparted by differential microtubule-binding affinities in the mother and daughter cells (see Fig. 7), and the likely role of phosphorylation in modulating the affinity of She1 for microtubules^13^, it’s also possible that a daughter cell-localized kinase phosphorylates She1 to reduce its microtubule binding in this compartment. Previous studies have identified the mitotic kinase Aurora B (Ipl1) and the mitogen-activated protein kinase Hog1 as candidates that may phosphorylate She1^13,37,64^. Although Ipl1 does not exhibit any known asymmetries in budding yeast, at least one upstream activator of Hog1 is indeed enriched in the daughter cell (Sho1)^65^, suggesting this kinase may be key in modulating She1 asymmetry, which is likely critical for the polarization of dynein-mediated spindle migration toward the daughter cell.

Although She1 is the only known MAP that has the capacity to polarize dynein-mediated cargo transport, recent studies have revealed several MAPs in higher eukaryotes that indeed impact motor-mediated cargo transport. For instance, MAP4 and MAP9 affect dynein-dynactin transport functions in higher eukaryotes^6–8^. Of note, depletion of MAP4 leads to hyperactive cortical dynein activity and pronounced spindle movements, much like deletion of She1 in budding yeast^8^. The mechanism by which MAP4 functions is unclear; however, MAP9 precludes the interaction between microtubules and dynein-dynactin complexes by blocking dynactin-microtubule binding^7^. Although there is limited sequence homology among the MAP family (restricted to the microtubule-binding domains), many of them are enriched with regions of intrinsic disorder (95%, 94%, and 96% of MAP9, MAP4 and She1, respectively, are predicted to be disordered according to MetaDisorder^66^). Thus, although sequence alignments reveal no clear She1 homolog in higher eukaryotes, MAP4 and/or MAP9 may have evolved from She1, or a common ancestor.

## ACKNOWLEDGEMENTS

We are extremely grateful to Luke Rice and members of his laboratory for sharing reagents (yeast strain and plasmids) and expertise pertaining to the expression, purification, and assembly of yeast tubulin. We are also grateful to Jeffrey Moore for sharing the *nip100^ΔCAP-gly^* yeast strain, and members of the Markus and DeLuca laboratories for valuable discussions. This work was funded by the NIH/NIGMS (R01GM118492 to S.M.M., and R01GM079373, P01GM105537, and R35GM134842 to C.L.A.). C.L. Asbury was also funded by the David and Lucile Packard Foundation (fellowship 2006-30521). M.E. Bailey was supported by an NIH Interdisciplinary Training Fellowship (T32CA080416).

## METHODS

### Plasmid generation

For expression and purification of She1-NLS^SV40^-HaloTag (see below), we introduced sequence encoding the nuclear localization sequence (NLS) from SV40 large T antigen (hereafter referred to as “NLS^SV40^”) between She1 and the C-terminal HaloTag. Briefly, the C-terminal HaloTag sequence was PCR amplified from pProEX-HTb-TEV:*SHE1-HALO*^13^ using a forward primer that includes a sequence that encodes a short linker (Gly-Ser-Gly-Ser) followed by the NLS^SV40^ (Ala-Ala-Ala-Pro-Lys-Lys-Lys-Arg-Lys-Val-Gly). This PCR product was assembled into pProEX-HTb-TEV:*SHE1-HALO* digested with BamHI and NotI (which excises the HaloTag) using Gibson assembly, yielding *pProEX-HTb-TEV:SHE1-NLS^SV40^-HALO*. Note this sequence (linker and NLS^SV40^) is identical to the one introduced at the 3’ end of the *SHE1* locus to generate *she1^NLS^*.

### Media and strain construction

Strains are derived from either YEF473A or W303, and are available upon request (listed in Supplementary Table 1). We transformed yeast strains using the lithium acetate method^67^. Engineered yeast strains (*e.g., she1^NLS^, dyn1m^MTBD-HA^*) were constructed by PCR product-mediated transformation^68^ or by mating followed by tetrad dissection. Proper tagging and mutagenesis was confirmed by PCR and/or by sequencing. Fluorescent tubulin-expressing yeast strains were generated using plasmids and strategies described previously^69,70^. Yeast synthetic defined (SD) media was obtained from Sunrise Science Products (San Diego, CA).

### Live-cell imaging experiments

Assessment of cell cycle progression and spindle positioning (see Figures 1 and S1) was performed by imaging cells in the CellAsic ONIX system using microfluidic cassettes designed for haploid yeast cells (Y04C; MilliporeSigma). In brief, after an overnight growth in SD complete media (supplemented with 2% glucose) at 30°C, cells were diluted 50-fold into the cell inlet well of the microfluidic cassette, which was primed with SD complete prior to addition of cells (per manufacturer’s instructions). Pressure was maintained at a 7.0 psi throughout the imaging period to ensure a constant replenishment of media into the cassette, which was set to 30°C. Note that temperature is not directly monitored by the CellAsic system, and is thus affected by ambient room temperature. To account for temperature differences between experiments, wild-type and mutant cells were imaged simultaneously by introducing respective cells into adjacent imaging chambers of the microfluidics cassette. Z-stacks (7 steps with 0.5 μm spacing) from multiple XY coordinates (10 for replicate 1, and 5 for replicate 2) were acquired for 10 hours at 90 second intervals. For spindle dynamics and nuclear translocation assays, mid-log phase cells were arrested with 200 mM hydroxyurea (HU) for 2.5 hours in SD complete supplemented with 2% glucose, and then applied to slide-mounted agarose ‘pads’ comprised of 1.7% agarose dissolved in SD complete supplemented with 2% glucose and 200 mM HU for confocal fluorescence microscopy. Full Z-stacks (19 planes with 0.2 μm spacing) of GFP-labeled microtubules (GFP-Tub1; for Figures 1G, 4, 5 and 6), and/or NLS-3mCherry (for Figure 3) were acquired every 10 seconds for 15 minutes on a stage pre-warmed to 28°C. To image She1-3GFP (for Figure 7), mid-log phase cells cultured in SD complete media supplemented with 2% glucose were pelleted, and then resuspended in SD complete media supplemented with 2% galactose (to activate the *GAL1* promoter). After 30 minutes, the cells were pelleted again, and resuspended in SD complete media supplemented with 2% galactose and 200 mM HU for 2.5 hours followed by applying them to slide-mounted agarose pads for confocal microscopy (for a total galactose induction time of 3 hours). Z-stacks (7 steps with 0.6 μm spacing) of mRuby2-Tub1 and She1-3GFP were acquired every 20 seconds for 1 minute.

Images were collected on a Nikon Ti-E microscope equipped with a 1.49 NA 100X TIRF objective, a Ti-S-E motorized stage, piezo Z-control (Physik Instrumente), an iXon DU888 cooled EM-CCD camera (Andor), a stage-top incubation system (Okolab), and a spinning disc confocal scanner unit (CSUX1; Yokogawa). 445 nm, 488 nm, 514 nm, 561 nm and 594 nm lasers housed in an LU-NV laser unit equipped with AOTF control (Nikon) were used to excite mTurquoise2, GFP, Venus, mRuby2 and mCherry, respectively, which were used with emission filters mounted in a filter wheel (ET480/40m for mTurquoise2, ET525/50M for GFP, ET520/40M for Venus, and ET632/60m for mRuby2 and mCherry; Chroma). The microscope was controlled with NIS Elements (Nikon).

For analysis of the cell cycle progression images, we used the following morphological features to define temporal landmarks in cell cycle progression: spindle pole body duplication was identified as the first frame when two SPBs could be spatially resolved; anaphase onset was defined as the first frame when the spindle began to elongate; cytokinesis was defined as the first frame when independent movement of the spindle pole bodies was apparent (in which they moved with respect to each other, indicating complete spindle disassembly).

### Protein purification

We purified She1-HaloTag as previously described^10^. Briefly, *E. coli* BL21 (Rosetta DE3 pLysS) cells transformed with pProEX-HTb-TEV:*SHE1-HALO* (or *pProEX-HTb-TEV:SHE1-NLS^SV40^-HALO*) were grown at 37°C in LB supplemented with 1% glucose, 100 μg/ml carbenicillin and 34 μg/ml chloramphenicol to OD_600_ 0.4-0.6, shifted to 16°C for 2 hours, then induced with 0.1 mM IPTG for 14-16 hours at 16°C. The cells were harvested, washed with cold water, and stored at −80°C. Cells were thawed in 0.5 volume of cold 2X lysis buffer [1X buffer: 30 mM HEPES pH 7.2, 50 mM potassium acetate, 2 mM magnesium acetate, 0.2 mM EGTA, 10% glycerol, 1 mM DTT, and protease inhibitor tablets (Pierce)] and then lysed by sonication (5 × 30 second pulses) with 1 minute on ice between each pulse. The lysate was clarified at 22,000 × g for 20 minutes, adjusted to 0.005% triton X-100, then incubated with glutathione agarose beads for 1 hour at 4°C. The resin was then washed three times in wash buffer (30 mM HEPES pH 7.2, 50 mM potassium acetate, 2 mM magnesium acetate, 0.2 mM EGTA, 300 mM KCl, 0.005% Triton X-100, 10% glycerol, 1 mM DTT, protease inhibitor tablets) and twice in TEV digest buffer (10 mM Tris pH 8.0, 150 mM KCl, 0.005% Triton X-100, 10% glycerol, 1 mM DTT). To fluorescently label She1-HALO, the bead-bound protein was incubated with 6.7 μM HaloTag-TMR ligand (Promega) for 15 minutes at room temperature. The resin was then washed three more times in TEV digest buffer to remove unbound ligand, then incubated in TEV buffer supplemented with TEV protease for 1 hour at 16°C. The resulting eluate was collected using a centrifugal filter unit (0.1 μm, Millipore), aliquoted, drop frozen in liquid nitrogen and stored at −80°C.

Purification of ZZ-TEV-6His-GFP-3HA-GST-dynein331-HALO (under the control of the galactose-inducible promoter, *GAL1p*) was performed as previously described^10,54^. Briefly, yeast cultures were grown in YPA supplemented with 2% galactose, harvested, washed with cold water, and then resuspended in a small volume of water. The resuspended cell pellet was drop frozen into liquid nitrogen and then lysed in a coffee grinder (Hamilton Beach). After lysis, 0.25 volume of 4X lysis buffer (1X buffer: 30 mM HEPES, pH 7.2, 50 mM potassium acetate, 2 mM magnesium acetate, 0.2 mM EGTA, 1 mM DTT, 0.1 mM Mg-ATP, 0.5 mM Pefabloc SC, 0.7 μg/ml Pepstatin) was added, and the lysate was clarified at 22,000 x g for 20 min. The supernatant was then bound to IgG sepharose 6 fast flow resin (GE) for 1 hour at 4°C, which was subsequently washed three times in wash buffer (30 mM HEPES, pH 7.2, 50 mM potassium acetate, 2 mM magnesium acetate, 0.2 mM EGTA, 300 mM KCl, 0.005% Triton X-100, 10% glycerol, 1 mM DTT, 0.1 mM Mg-ATP, 0.5 mM Pefabloc SC, 0.7 μg/ml Pepstatin), and twice in TEV buffer (50 mM Tris, pH 8.0, 150 mM potassium acetate, 2 mM magnesium acetate, 1 mM EGTA, 0.005% Triton X-100, 10% glycerol, 1 mM DTT, 0.1 mM Mg-ATP, 0.5 mM Pefabloc SC). To fluorescently label 6His-GFP-GST-3HA-dynein331-HALO (for single molecule analyses), the bead-bound protein was incubated with either 6.7 μM HaloTag-TMR or HaloTag-PEG-biotin ligand (Promega) for 15 minutes at room temperature. The resin was then washed four more times in TEV digest buffer, then incubated in TEV buffer supplemented with TEV protease for 1 hour. Following TEV digest, the bead solution was transferred to a spin column (Millipore) and centrifuged at 20,000 x g for 10 seconds. The resulting protein solution was aliquoted, flash frozen in liquid nitrogen, and then stored at −80°C.

Purification of yeast tubulin was performed essentially as described^71^ with minor modifications. Yeast cells (JEL1) co-transformed with p426Gal1:Tub1 and p424:Tub2-6His were grown in 50 ml of selective SD complete media (lacking uracil and tryptophan) supplemented with 2% glucose, and then transferred to 1 L of nonselective YPGL (2% peptone, 1% yeast extract, 3% glycerol, and 2% lactate; note, we grew 16 L of cells for a typical prep). When the cell density reached an OD600 between 5-9, 20 grams of galactose powder was added per liter of YPGL, and after 5 hours, cells were harvested, washed with water, and stored at −80°C. Approximately 75 gram of cells were thawed and resuspended in 70 ml of lysis buffer (50 mM HEPES pH 7.4, 500 mM NaCl, 10 mM MgSO_4_, 30 mM imidazole) supplemented with 50 μM GTP and cOmplete protease inhibitor cocktail (Roche), and lysed by 5-6 passes through a microfluidizer (LM10; Microfluidics) at 23,000 PSI, with 5 minutes on ice between each pass. After clarification (at 13,000 rpm for 30 minutes at 4°C), the supernatant was applied to a 5 ml HisTrap Ni-NTA column (GE) pre-equilibrated with 10 column volumes (CVs) of lysis buffer supplemented with GTP using an AKTA FPLC (GE). After washing the column with 10 CVs of lysis buffer supplemented with 50 μM GTP and 10 CVs of nickel wash buffer (25 mM PIPES pH 6.9, 1 mM MgSO_4_, 30 mM imidazole) supplemented with 50 μM GTP, bound protein was eluted with 6 CVs of elution buffer (25 mM PIPES pH 6.9, 1 mM MgSO_4_, 300 mM imidazole) supplemented with 50 μM GTP. Peak fractions (determined by absorbance at 260 nm) were pooled and treated with nuclease (Pierce Universal Nuclease; catalog #88702; 10 μl per 20 ml of eluate) for 15 minutes at room temperature, and then diluted with MonoQ buffer A (25 mM PIPES pH 6.9, 2 mM MgSO_4_, 1 mM EGTA) supplemented with 50 μM GTP such that the final imidazole concentration was 100 mM. The protein was then loaded onto a MonoQ 10/100GL anion exchange column pre-equilibrated with 5 CVs of 90% MonoQ buffer A (see above) and 10% MonoQ buffer B (25 mM PIPES pH 6.9, 2 mM MgSO_4_, 1 mM EGTA, 1 M NaCl) supplemented with 50 μM GTP, after which bound protein was eluted using with a 10-70% MonoQ buffer B gradient over 50 CVs. Peak tubulin fractions (determined by absorbance at 260 nm and SDS-PAGE) were pooled and concentrated (Amicon Ultra-4 30K; catalog #UFC803024) to 2.8 μM (concentration and aggregation were closely monitored using absorbance at 260 nm and 280 nm), and then dialyzed against tubulin storage buffer (10 mM PIPES pH 6.9, 1 mM MgSO_4_, 1 mM EGTA) supplemented with 50 μM GTP (Thermo Slide-A-Lyzer; catalog #66810). Resulting protein was aliquoted (50 μl), snap frozen in liquid nitrogen, and stored at −80°C.

### Optical Trapping

Anti-His-coated 0.44 μm microbeads (PSS4; Spherotech; prepared as described previously^72^) were incubated with purified 6His-GFP-3HA-GST-dynein_331_-HALO in dynein trapping buffer (30 mM HEPES pH 7.2, 2 mM Mg-Acetate, 1 mM EGTA) for 1 hour at 4°C. During the incubation, flow chambers (assembled from a glass slide, coverslip, and double stick tape) were prepared by sequential addition and incubation with the following solutions: (1) 1 mg/ml biotinylated BSA (Vector Labs #B-2007), (2) BRB80 (80 mM PIPES pH 6.9, 1 mM MgCl_2_, 1 mM EGTA, pH 6.9), (3) 0.33 mg/ml Avidin (Vector Labs #A-3100), (4) BRB80, and (5) GMPCPP-stabilized biotinylated microtubules (diluted in BRB80)^73^. After a 10 minute incubation, the chamber was washed with BRB80, and the dynein-coated microbeads diluted in trapping buffer supplemented with 1 mg/ml κ-casein, 8 mg/ml BSA, 1 mM DTT, 0.8 mM ATP, 4 mM MgSO_4_), 4.5 mg/ml glucose, 250 μg/ml glucose oxidase, 30 μg/ml catalase, were introduced into the chamber, which was then sealed with nail polish, and immediately used for data collection.

The optical trap was essentially as described^74^, and was operated in stationary mode, without feedback control (*i.e*., in “open loop” mode). Bead-trap separation was saved at 200 Hz and converted into force by multiplying by the trap stiffness, which ranged between 0.025 and 0.045 pN/nm. Custom analysis software written in Igor Pro (Wavemetrics) was used to estimate pre-stall speeds, stall forces, and stall times for individual bead motility events. Briefly, we defined the start of an event as the time at which the bead first moved beyond 3× the root-mean-square baseline noise. The end of an event was clearly identifiable as the time when the bead detached from the microtubule. The onset of stalling was chosen as the time at which the bead velocity, averaged over a sliding 2.5-s window, first fell below 2 nm/s. Pre-stall speed was then defined as the slope of a line fit to all the data between the start of the event and the onset of stalling and stall force was defined as the average force level during the stall event.

### In vitro microtubule-binding assays

We used total internal reflection fluorescence (TIRF) microscopy-based microtubule-binding assays to measure the binding affinity of She1 and She1^NLS^ for yeast microtubules. To prepare microtubules, 15 μl of tubulin polymerization buffer (500 mM PIPES pH 6.9, 5 mM MgSO_4_, 25% glycerol) along with epothilone B (50 μM final) and GTP (2 mM final) were added to one 50 μl aliquot of yeast tubulin (see above), which was then incubated at 30°C overnight. Flow chambers constructed from slides, double-stick tape, and plasma cleaned and silanized coverslips were coated with anti-His antibody (100 μg/ml, sc-8036, Santa Cruz), then blocked with 1% Pluronic F-127, after which microtubules (diluted to 0.75 μM in tubulin polymerization buffer supplemented with 50 μM epothilone B and 2 mM GTP) were added. After unbound microtubules were removed by washing with chamber wash buffer (a 9:1 mixture of the following two buffers: dynein assay buffer: 30 mM HEPES pH 7.2, 50 mM potassium acetate, 2 mM magnesium acetate, 0.2 mM EGTA, 10% glycerol, supplemented with 1 mM DTT, 50 μM epothilone B, and 2 mM GTP; and TEV buffer, as described above; this mixture was used to account for storage of purified She1-TMR in TEV buffer), purified She1-TMR (wild-type or She1^NLS^, diluted such that the final 9:1 buffer mixture described above was achieved for all concentrations of She1 and She1^NLS^) was introduced into the chamber. Images were collected on a Nikon Ti-E microscope (controlled with NIS Elements) equipped with a 1.49 NA 100X TIRF objective, a motorized stage, piezo Z-control (Physik Instrumente), and an iXon X3 DU897 cooled EM-CCD camera (Andor). A 561 nm laser (Coherent) was used along with a multi-pass quad filter cube set (C-TIRF for 405/488/561/638 nm; Chroma) and emission filters mounted in a filter wheel (525/50 nm, 600/50 nm, and 700/75 nm; Chroma) to image She1-TMR. To image non-fluorescent yeast microtubules, we used interference reflection microscopy (IRM), as previously described^75^. Excitation light for IRM was provided by a Sola SE light engine (Lumencor). To measure the degree of She1-microtubule binding, background-subtracted fluorescence intensities of She1-TMR were determined. Note that we accounted for potential differences in the extent of HaloTag labeling between She1 and She1^NLS^ by measuring fluorescence band intensities following imaging of acrylamide gels with each on a Typhoon gel imaging system (FLA 9500). Both absolute concentration and relative degree of HaloTag labeling were taken into account when calculating binding affinities. Binding curves and curve fitting for dissociation constants (where appropriate) were generated using GraphPad Prism.

### Spindle tracking and statistical analysis

Spindle tracking was performed on maximum intensity projections (XY) using a custom written MATLAB routine (as described previously^76^). Dynein-mediated spindle movements were manually selected from the tracking data to obtain the various metrics described in Figures 4B – E, and 5B – E. To determine the fraction of time the spindle centroid resides within 1 μm of the cell cortex (Figures 4F and 5F), an additional MATLAB routine was generated with which the user manually defines the cell cortex of the mother and bud cell. To determine the fraction of time the spindle resides within mother and daughter cell (Figures 4G and 5G), and the relative distance from the bud neck (Figures 4H and 5H), another MATLAB routine was generated in which the user manually defines the bud neck. For this latter routine, the edges of the cell were defined by cropping individual cell images such that the extreme left and right cell edges coincided with the cropped image.

P values were calculated using either an unpaired two-tailed Welch’s t-test, or the Mann-Whitney test, both of which were performed using GraphPad Prism. These tests were selected as follows: the unpaired two-tailed Welch’s t-test was used when the datasets in question were both determined to be normal (by the D’Agostino & Pearson test for normality; P > 0.05); in the case where only one (or neither) of the datasets were determined to be normal (P < 0.05), the Mann-Whiney test was used.

#### Cell lysis and immunoblotting

For Western blotting, yeast cultures were grown at 30°C in 3 ml SD supplemented with 2% glucose and harvested. Equal numbers of cells were pelleted and resuspended in 0.2 ml of 0.1 M NaOH and incubated for 10 min at room temperature as described^77^. After centrifugation, the resulting cell pellet was resuspended in sample buffer and heated to 100°C for 3 min. Lysates were separated on an SDS polyacrylamide gel and electroblotted to PVDF in 25 mM Tris and 192 mM glycine supplemented with 0.05% SDS and 10% methanol for 30 min. Rabbit anti–c-Myc polyclonal (GenScript) and HRP-conjugated goat anti-rabbit antibody (Jackson ImmunoResearch Laboratories) were used at 1:1000, and 1:3000, respectively. Total protein (using Stain-Free technology; BioRad) and chemiluminescence signal were both acquired on a BioRad ChemiDoc MP gel documentation system without saturating the camera’s pixels. Band intensities (and background values) were measured using NIH ImageJ.

## SUPPLEMENTARY FIGURE LEGENDS AND TABLE

**Figure S1.**
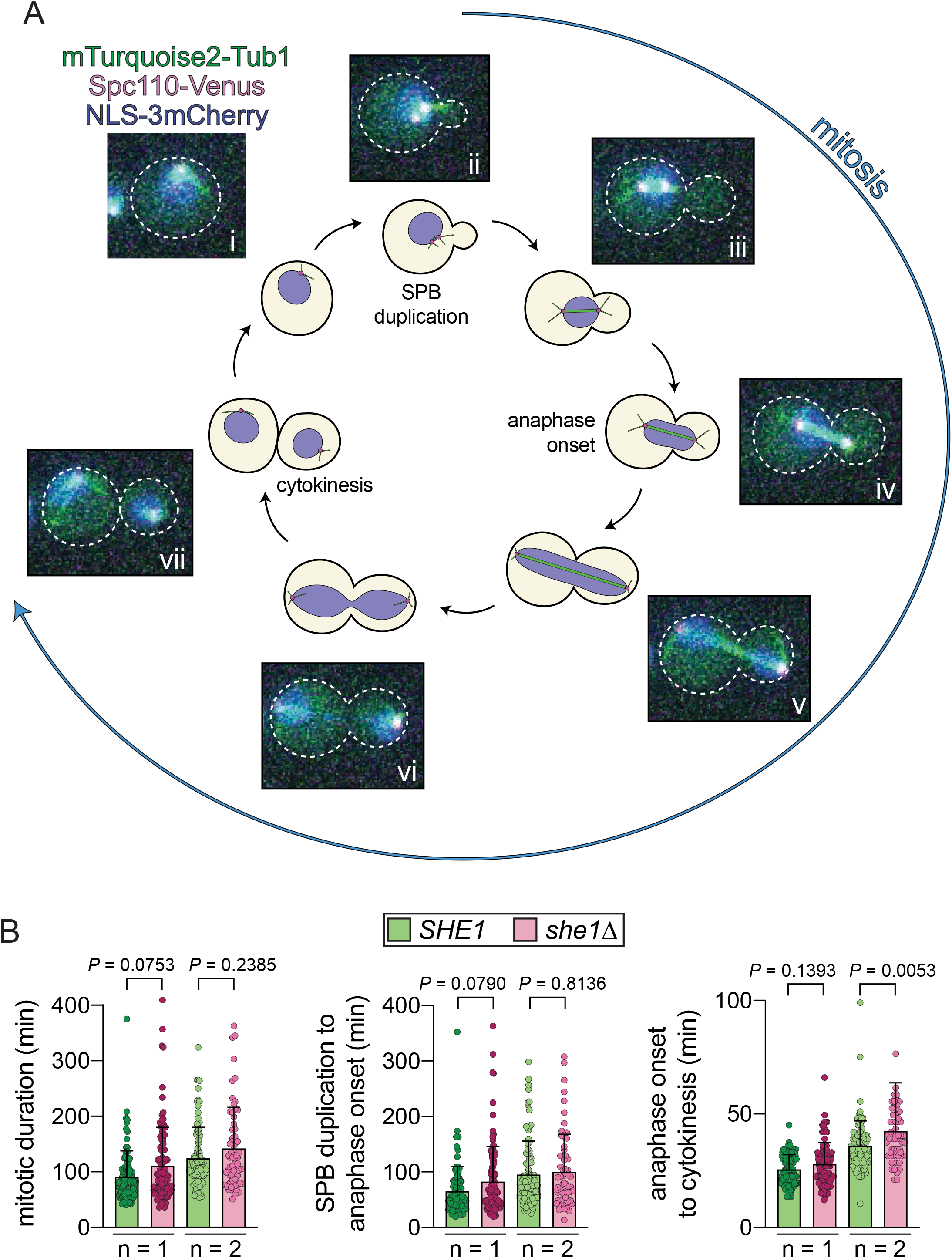
Cell cycle progression analysis. (A) Representative time-lapse confocal fluorescence microscopy images of a cell expressing mTurquoise2-Tub1 (to visualize microtubules), Spc110-Venus (to visualize spindle pole bodies, or SPBs), and NLS-3mCherry (to visualize the nucleus). Cells were grown in a microfluidics cassette (CellAsic ONIX; see Methods), and imaged over the course of several cell cycles. (B) Plots depicting absolute time intervals between indicated temporal landmarks for independent replicates 1 and 2 (see Figure 1 for normalized data, and n values). Images for independent replicates for *SHE1* and *she1*Δ cells were acquired simultaneously (*i.e*., *SHE1* and *she1*Δ replicate #1 were acquired together; see Methods) to account for potential differences in room temperature.

**Figure S2.**
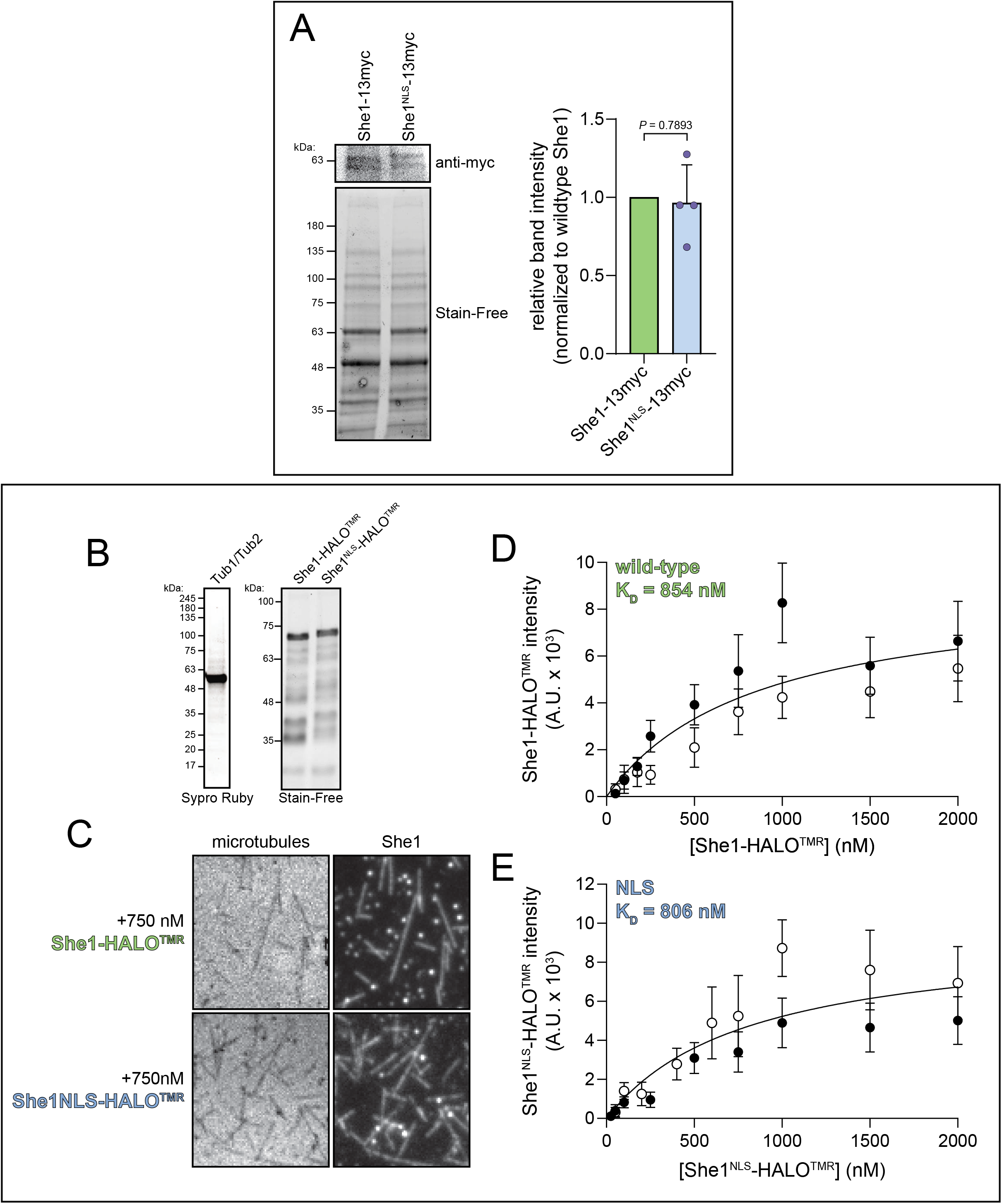
Addition of a C-terminal nuclear localization signal peptide to She1 does not disrupt its expression or microtubule-binding activity. (A) Representative immunoblot (top) and Stain-Free image (bottom) of extracts prepared from cells expressing She1-13myc (wild-type or She1^NLS^) and separated by SDS PAGE. Total protein was imaged using Stain-Free technology (BioRad; bottom), and used as a loading control. Background-subtracted band intensities (corrected for differences in sample loading) for She1^NLS^-13myc from each replicate (n = 2 independent experiments, with 2 lysate concentrations per immunoblot) were directly compared to that from each respective wild-type, which was normalized to 1. (B) Recombinant proteins used in binding assays (left, alpha/beta-tubulin from yeast; right, She1 proteins from bacteria). (C) Representative images of microtubule-bound She1 (wild-type and She1^NLS^) used in quantitation of binding affinities. (D and E) Plots depicting relative microtubule binding of wild-type (D) and She1^NLS^ (E) as a function of concentration (open and closed circles represent values obtained from independent replicates; n = 2). Background subtracted intensity values for microtubule-bound She1 were plotted against concentration, and the data were fit (using GraphPad Prism) to obtain K_D_ values, as shown.

**Figure S3.**
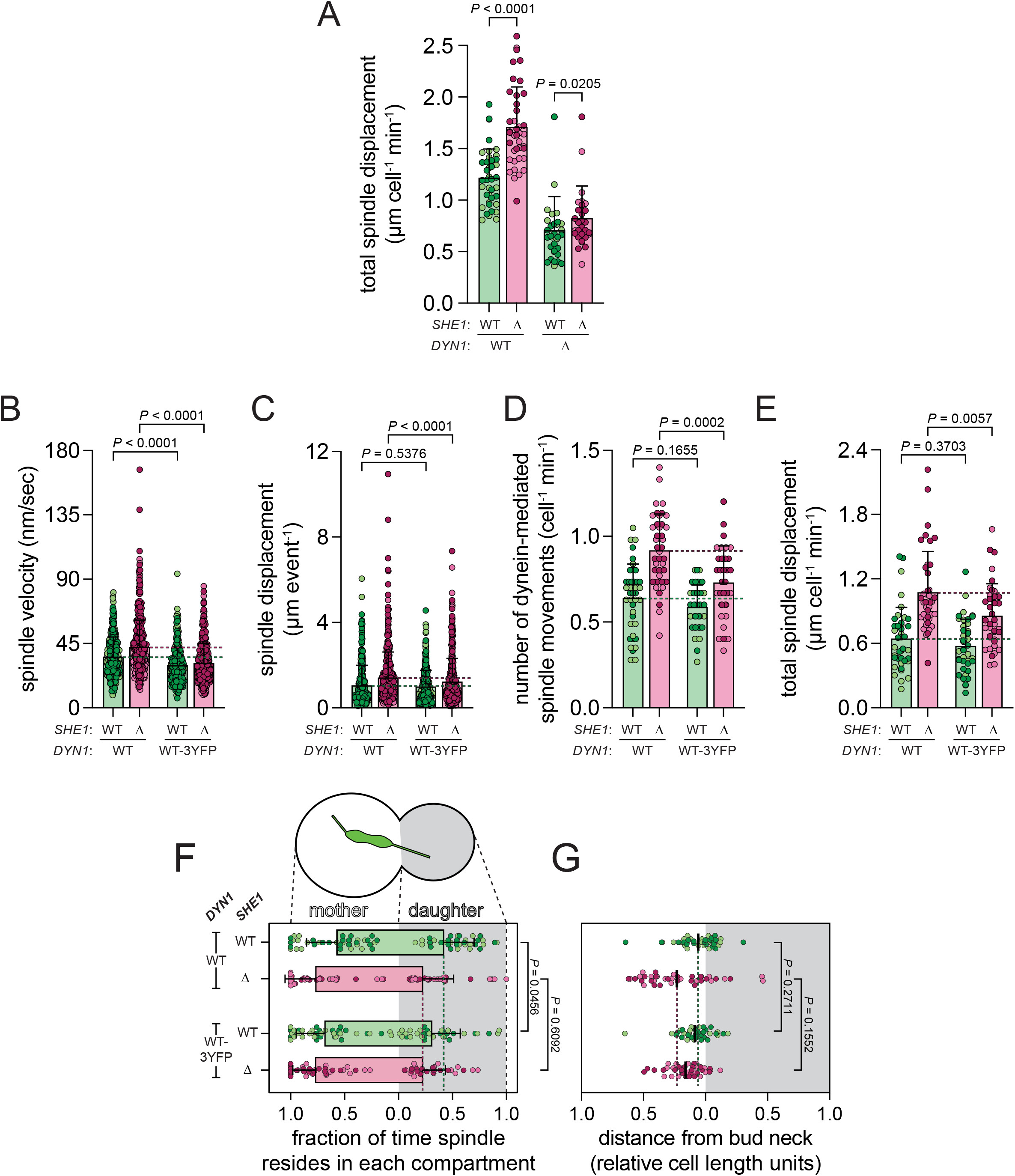
Dynein and She1 are required for robust spindle movements, and C-terminal 3YFP tag on Dyn1 attenuates she1Δ-dependent phenotypes. (A) Plots depicting total spindle displacement per minute for indicated yeast strains (from all tracking data, not just dynein-mediated displacement; from left to right, n = 40, 40, 35 and 35 cells from 2 independent replicates). (B – E) Dynein-mediated spindle movements were manually selected from the tracking data, from which velocity (B), displacement (C, per event; or, E, per minute), and the number of dynein-mediated spindle movements per minute (D) were obtained (mean ± standard deviation is overlaid with scatter plot of all data points; from left to right). See Figures 4 and 5 for n values. P values were calculated using either the Mann-Whitney or an unpaired twotailed Welch’s t-test (see Methods). (F and G) Plots depicting the relative number of spindle coordinates that reside within the mother and daughter cell (F; circles represent fraction of time spindle resides within each compartment for individual cells), and, the mean position of the spindle centroid (along the longitudinal mother-daughter axis only) for each cell (G; circles represent mean spindle position for individual cells over the course of a 15 minute movie, and lines indicate mean values for all cells). See Figures 4 and 5 for n values. P values were calculated using the Mann-Whitney test (see Methods). For all panels, light and dark color hues indicate data points from independent replicates (“WT-3YFP”, *DYN1-3YFP*). For panels B – G, green and red dashed lines delineate mean values for *DYN1* and *DYN1 she1*Δ cells, respectively. See Figures 4 and 5 for P values comparing *SHE1* datasets to those from *she1*Δ cells.

**Figure S4.**
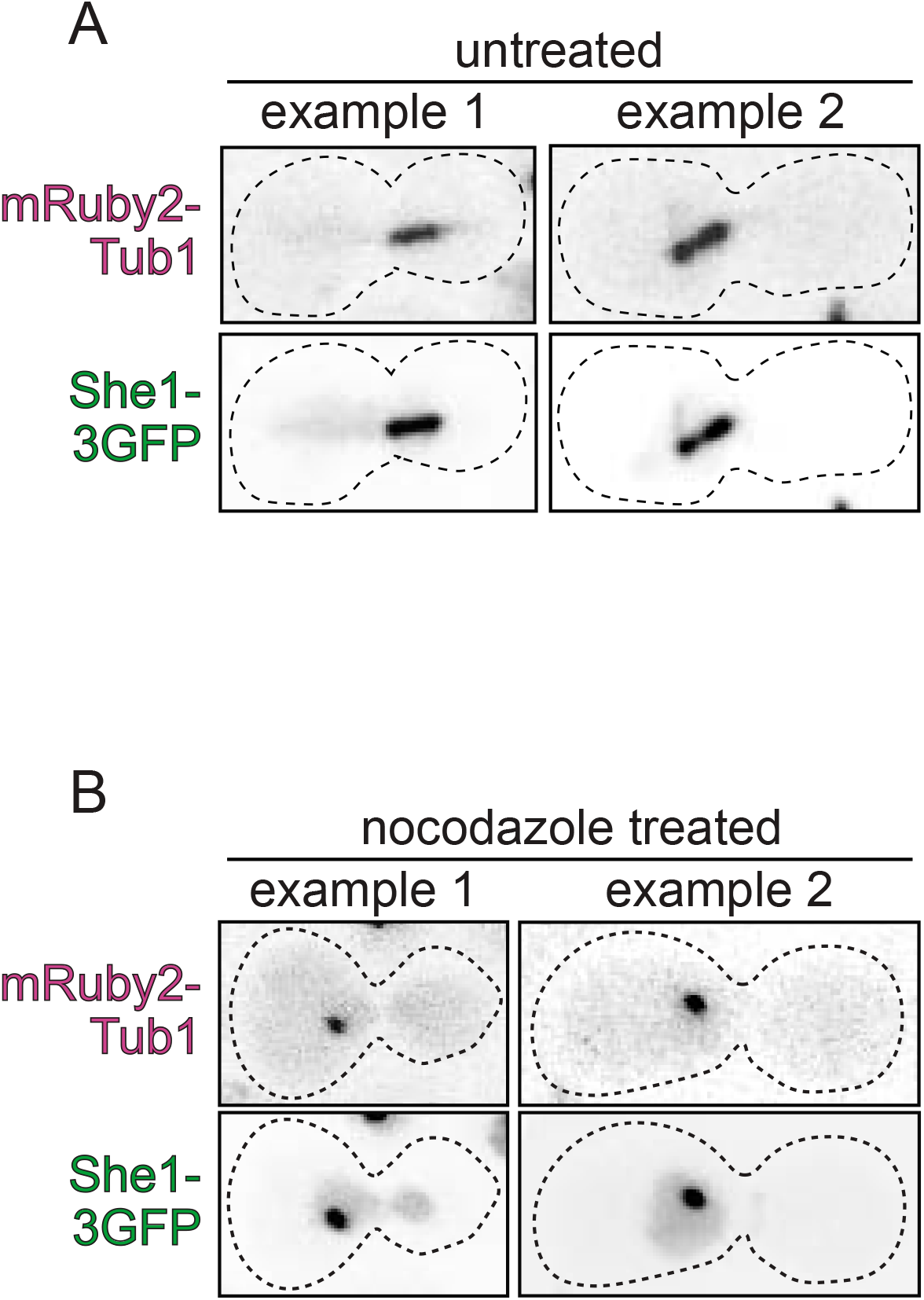
Treatment with nocodazole eliminates astral and spindle microtubules. (A and B) Representative microscopy images of hydroxyurea-arrested cells expressing mRuby2-Tub1 and overexpressing She1-3GFP (as described in Figure 7). Those in panels A and B are examples of untreated cells, and those treated with 100 μM nocodazole for 30 minutes prior to imaging, respectively. Note the absence of clear microtubule-based structures (astral and spindle microtubules). However, we noted the appearance of bright spots in both mRuby2-Tub1 and She1-3GFP channels upon nocodazole treatment, which may be a collapsed spindle and/or single SPB with very short microtubules to which She1 could bind.

**Table S1.**
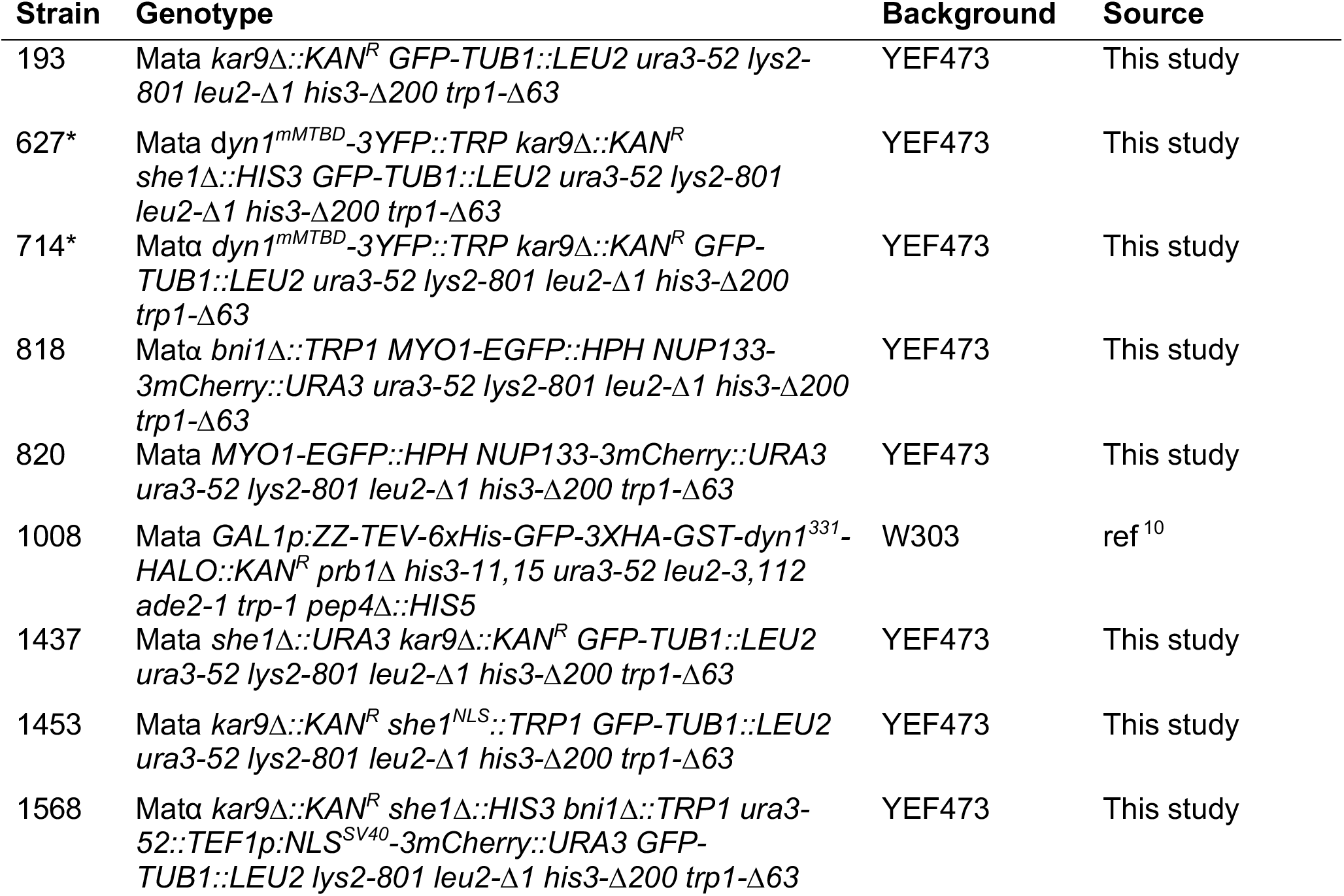
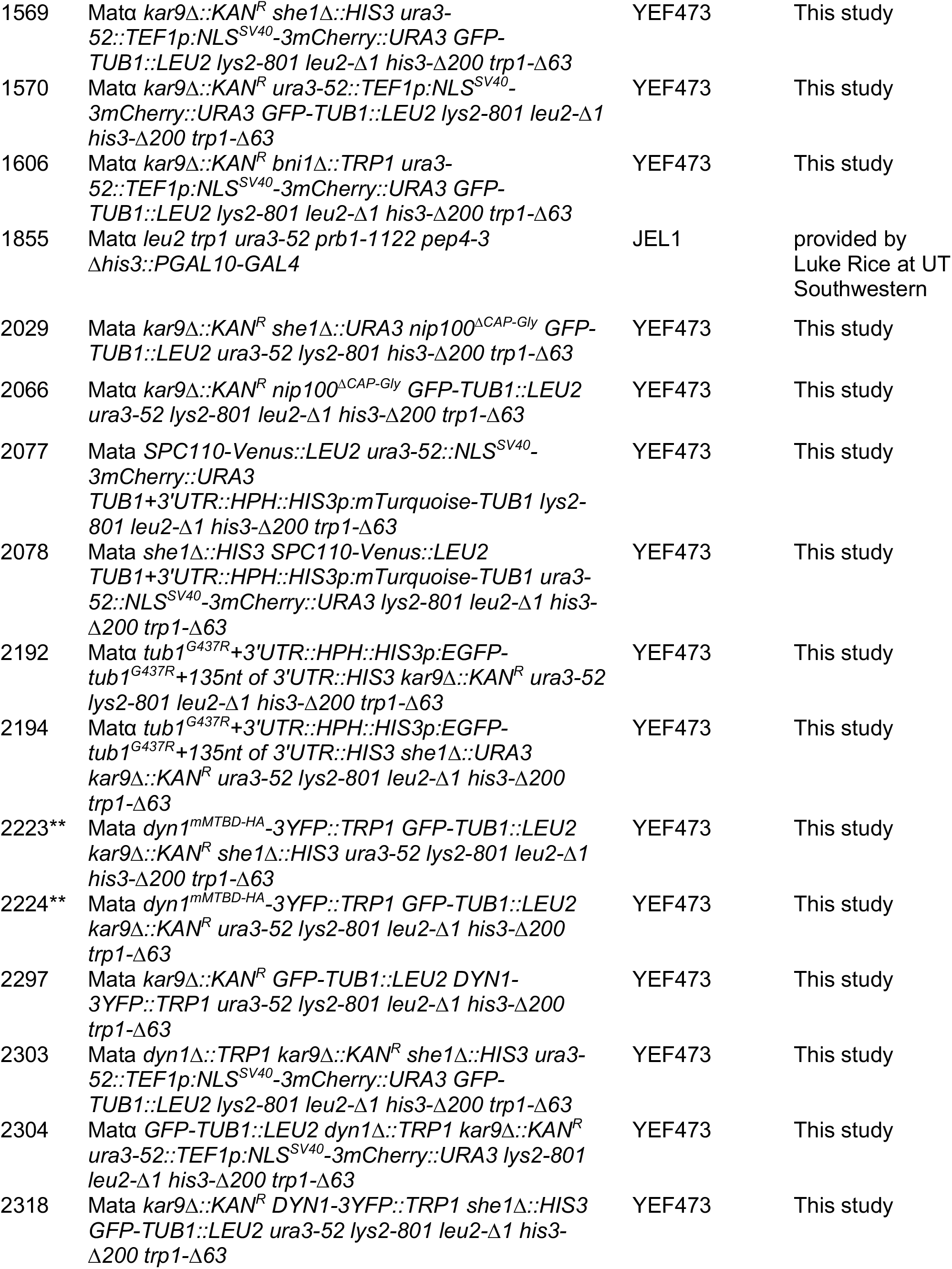
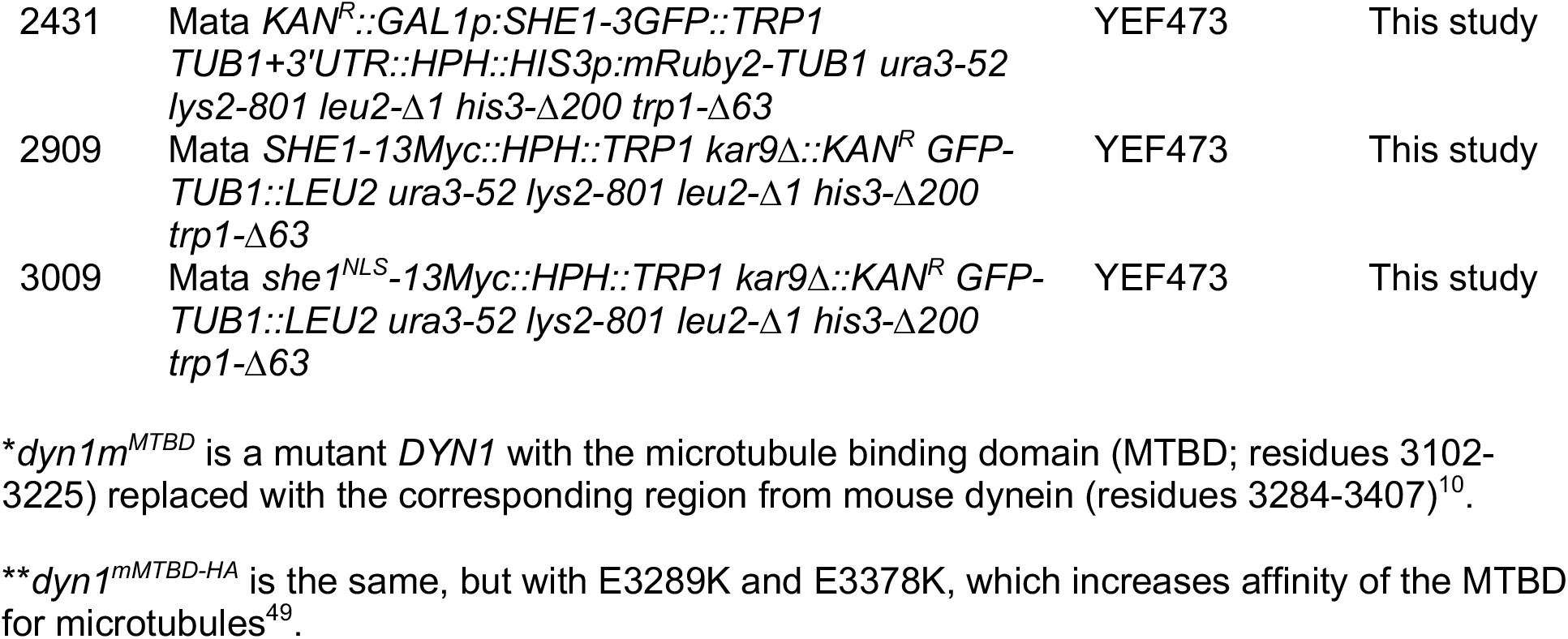
List of yeast strains used throughout the study.

